# Conditional language models enable the efficient design of proficient enzymes

**DOI:** 10.1101/2024.05.03.592223

**Authors:** Geraldene Munsamy, Ramiro Illanes-Vicioso, Silvia Funcillo, Ioanna T. Nakou, Sebastian Lindner, Gavin Ayres, Lesley S. Sheehan, Steven Moss, Ulrich Eckhard, Philipp Lorenz, Noelia Ferruz

## Abstract

The design of functional enzymes holds promise for transformative solutions across various domains but presents significant challenges. Inspired by the success of language models in generating nature-like proteins, we explored the potential of an enzyme-specific language model in designing catalytically active artificial enzymes. Here, we introduce ZymCTRL (’enzyme control’), a conditional language model trained on the enzyme sequence space, capable of generating enzymes based on user-defined specifications. Experimental validation at diverse data regimes and for different enzyme families demonstrated ZymCTRL’s ability to generate active enzymes across various sequence identity ranges. Specifically, we describe the design of carbonic anhydrases and lactate dehydrogenases in zero-shot, without requiring further training of the model, and showcasing activity at sequence identities below 40% compared to natural proteins. Biophysical analysis confirmed the globularity and well-folded nature of the generated sequences. Furthermore, fine-tuning the model enabled the generation of lactate dehydrogenases outside of natural sequence space but with activity comparable to their natural counterparts. Two of the artificial lactate dehydrogenases were selected for scale production and successfully lyophilised, maintaining activity and demonstrating preliminary conversion in one-pot enzymatic cascades under extreme conditions. Our findings open a new door towards the rapid and cost-effective design of artificial proficient enzymes. The model and dataset are freely available to the community.

## Introduction

Enzymes are captivating nanomachines with the ability to accelerate chemical transformations by several orders of magnitude. From converting atmospheric CO_2_ into valuable chemicals^1^ to facilitating the production of light^2^, natural enzymes exhibit a diverse array of functionalities, all achieved under mild conditions without the use of toxic solvents^3^. This inherent versatility has positioned enzymes as highly sought-after materials, promising to establish greener chemistries for myriad applications and reduce costs^4^. Indeed, significant advances have been made in the past two decades, including augmenting the thermostability and activity of enzymes like PETase^5^ and engineering *de novo* enzymatic reactions, including the Kemp elimination^6^, Diels-Alder^7^, retro-aldol^8^, and Morita-Baylis-Hilman^9^. Despite this enormous potential, it is evident that we still fail to design enzymes as proficient as natural ones, with labour-intensive processes that can span several years and involve various rounds of evolution^10^.

The reasons for these challenges are varied, including our incomplete understanding of the intricate molecular processes governing biochemical interactions and catalysis, the costly nature of the experimental assays, and the astronomical space of possible enzyme sequences - with most variants leading to loss of stability or function^11^. Artificial intelligence (AI) methods are emerging as powerful tools that outperform previous results across all disciplines. In the protein research realm, they tackle some of the previous limitations effectively and have unquestionably promoted a paradigm shift in the field^12^. Among AI models, Large language models (LLMs) are manifesting unprecedented performance, as evidenced by the recent ChatGPT or Gemini agents^13^. Protein language models (pLMs) have likewise showcased impressive success, including predicting the effect of mutations^14^ and protein structures^15,16^ by learning statistics of coevolving residues^17^ and allowing the step-wise design of *de novo* proteins^18^. In the context of enzyme design, pLMs and other generative models have provided artificial variants for several enzyme families^19,20^, including artificial TEM-1 β-lactamases^21^, lysozymes^22^, luciferases^2^, malate dehydrogenases^23^, superoxide dismutases^24^, chorismate mutases^25^, and CRISPR-Cas genome editors^26^ in the last three years alone^27^. While these works highlight the potential of AI architectures in the protein realm, the implementation of a model that produces highly active enzymes without the need for further training and with high success rates remains a longstanding goal in the field.

One limitation of most autoregressive pLMs is that these exert limited control over the properties of generated sequences in zero-shot scenarios, i.e. when generating sequences without additional training. Techniques such as fine-tuning on specific families^22^, prompt engineering^28^, or high-throughput sequence generation followed by property filtering have been employed to address this limitation^12^. Nevertheless, the ideal solution would be an end-to-end model capable of generating sequences based on user-defined prompts. This concept has been realised in models like CTRL^29^, which utilises control tags to guide text generation, and Progen^30^, a pLM trained with the potential to generate upon labels defining biological processes, cellular components, function, or taxonomy, recently resulting in the production of active lysosomes after fine-tuning^22^.

Here, we hypothesised that a conditional, autoregressive pLM trained on the known enzyme sequence space could learn an internal grammar of enzymatic activity and generate active enzymes upon user-defined requests. To test this hypothesis, we developed ZymCTRL (short for enzyme control), a conditional pLM trained to generate enzymes based on specific catalytic reactions. ZymCTRL was trained on the publicly available Uniprot database, comprising 37M enzyme sequences annotated with enzyme commission (EC) numbers at the time of training^31^. Each sequence was linked to its associated EC class during training, enabling the model to learn sequence features specific to each catalytic reaction. To address the known issue of lack of representativeness in certain families, we tokenised EC classes, allowing the model to transfer learned insights across catalytic reactions.

We experimentally test the performance of the model under different scenarios. First, we examine 20 beta carbonic anhydrases generated in zero-shot, with identities below 50% to the natural space. Carbonic anhydrases are the fastest enzymes known in nature, with catalytic efficiencies approaching the diffusion limit^32^. Therefore, this enzyme family constitutes a very astringent test of ZymCTRL capabilities. Seven of the artificial carbonic anhydrases showed activities, with two close to natural ones, with sequence identities in the range of 35 - 50%. Second, in order to address potential biases in pLMs due to unequal sequence sampling across the tree of life in public databases^33^, we fine-tuned ZymCTRL on a diverse set of metagenomic lactate dehydrogenase (LDH) sequences derived from Basecamp Research’s internal graph database. We show that LDH sequences generated after fine-tuning are more likely to pass *in silico* quality metrics than zero-shot generated sequences. 20 generated sequences were selected for testing (ten zero-shot and ten fine-tuned), all of which expressed and a majority of which showed activity in line with the natural controls. Two were selected for further studies, scaled up and lyophilised, retaining activity and showing mild conversion in one-pot enzymatic cascades even at extreme pH and temperature conditions, highlighting the usability of the generated sequences in industrial contexts. Given the ability of enzymes to catalyse industrial processes in an environmentally friendly manner, we believe that ZymCTRL represents a timely advancement toward conditional, cost-effective enzyme design. To benefit the scientific community, we have made this model freely available at https://huggingface.co/AI4PD/ZymCTRL.

### Training ZymCTRL: A model conditioned on enzyme classes

We have trained ZymCTRL, a conditional pLM capable of generating enzyme sequences that fulfil a user-defined catalytic reaction (**Fig. 1a, b**). Our objective was to train a foundation model that can generate high-quality enzymes conditioned on a specific enzyme class without needing further training, providing complete control to the end users. To this end, we used the Transformer architecture’s decoder module^34^ and trained it with an autoregressive objective on the Uniprot database^31^ to give a 738 M parameter-sized model (**Methods**). During training, each sequence was passed along with its corresponding EC class, ensuring that ZymCTRL learned a joint distribution of sequence-function relationships.

**Figure 1:**
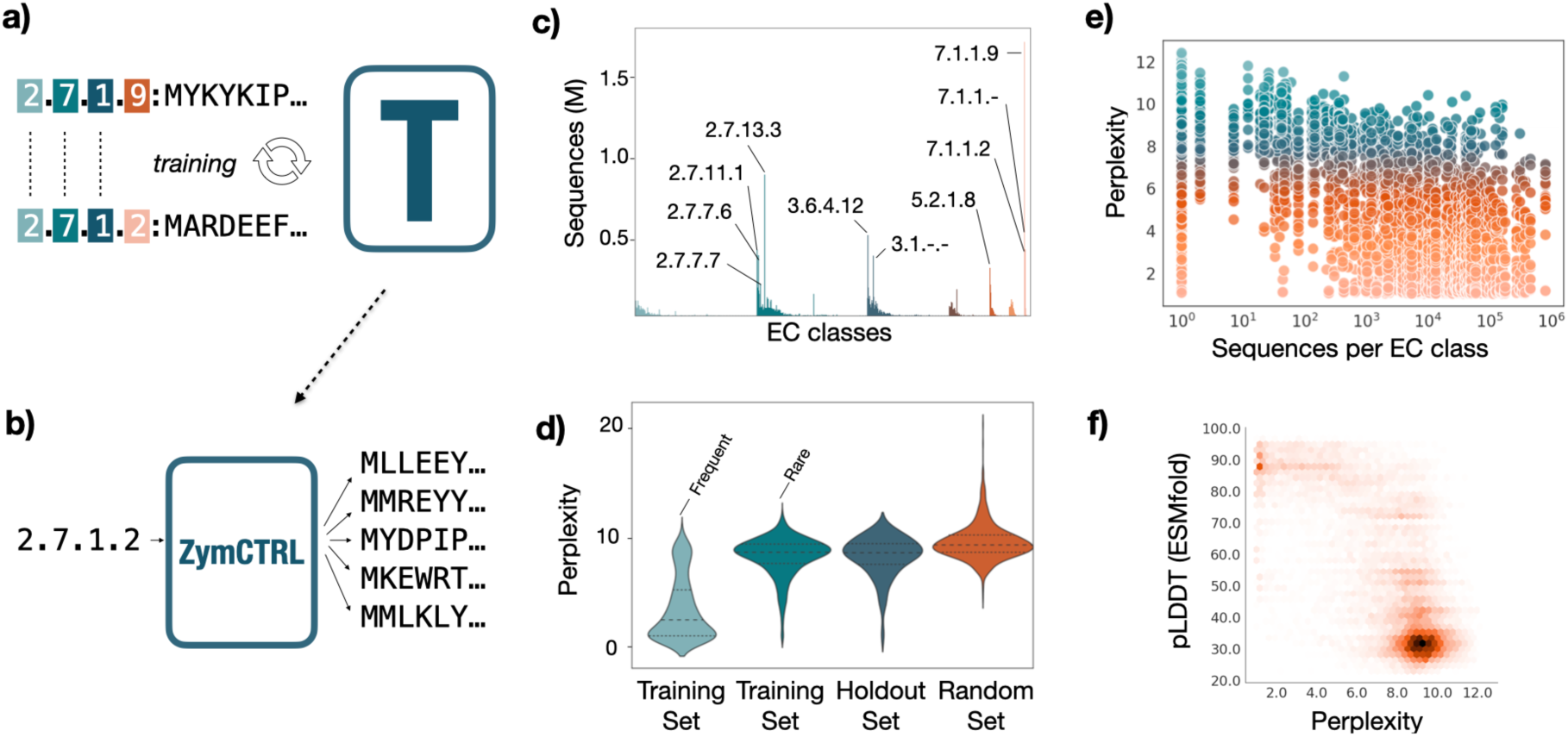
Training and generation process and sequences in the training database. **(a)** During pre-training, a decoder-only Transformer (T) model learns the relationship among sequences and their tokenised labels in an iterative process. The EC classes are also tokenised at the character level, allowing the model to infer relationships among the EC groups. **(b)** At inference time, users can specify a target catalytic reaction as a condition for the model’s generation, such as ‘2.7.1.2’: ‘glucokinases’. **(c)** Sequences in the database classified by EC number, labelled are the ten largest classes. **(d)** Perplexity for different groups of generated sequences. **(e)** Perplexity as a function of the number of sequences per EC class in the training set. **(f)** pLDDT values obtained with ESMFold as a function of perplexity, suggesting low-perplexity sequences are more prone to lead to ordered structures.

EC numbers feature a four-level hierarchy, with each successive number defining the catalytic activity more precisely. For example, enzymes classified as EC: 2.1.1.13 are transferases (first level), transferring one-carbon groups (second level), such as methyl (third level), to specifically regenerate methionine from homocysteine (fourth level). The annotated enzymes, however, feature large imbalances among classes, with some being significantly more populated than others: while the top 100 most populated classes encompass 37% of the sequences in the dataset, 9% of the classes only include one sequence (**Fig. 1c**).

There are a few possibilities to partly alleviate representation bias during training. One strategy would be to ensure an equivalent number of members per class. While this approach would deplete biases, it would also not exploit a significant proportion of this valuable annotated data. In contrast, we envisioned a training strategy where the model could transfer knowledge from populated to underrepresented classes. In particular, we tokenised the EC labels by subclasses (**Methods**) to promote that the model infers patterns within the same group (e.g., EC classes 2.7.1.9 and 2.7.1.2) and to understand the minimal requisites for good-quality per class generation (**Fig. 1a, b**).

In this sense, we analysed the perplexities for sequences in rare EC classes, i.e., EC classes with only one sequence in the training set. Perplexity is a measure of the model’s understanding of a given sequence and is mathematically defined as the exponential of the negative log-likelihood of the sequence (**Methods**). More intuitively, lower values indicate sequences for which the model has higher confidence. While sequences generated from EC classes with abundant data points exhibit lower perplexities, sequences from rare EC classes produce average perplexities lower than those from holdout and implausible (random) EC labels (**Fig. 1d, Table S1**). We indeed observe several sequences with very low perplexities in the case of EC classes with less than ten sequences, suggesting data transfer among classes (**Fig. 1e**). We wondered whether lower perplexity values correlate with other assessments, and observed a strong correlation with ESMFold^15^ pLDDT values (**Fig. 1f**). Taken together, the results point out that the model confidently generates sequences with high pLDDTs at all data regimes, even for highly unrepresented families with a single member.

### The generated enzymes are novel yet predicted ordered and functional

One critical property of language models is that they can generalise on the training set and infer novel, unseen, yet coherent texts. This is a particularly interesting property for protein design since it provides the means to explore novel distant regions in sequence space, enabling the potential to design new functionalities and enhancing our understanding of protein function. To understand the extent to which ZymCTRL explores novel sequences, we ran MMseqs2^4^ searches on generated sequences versus the training set (Methods, **Fig. S1**, **Fig. 2a**).

**Figure 2:**
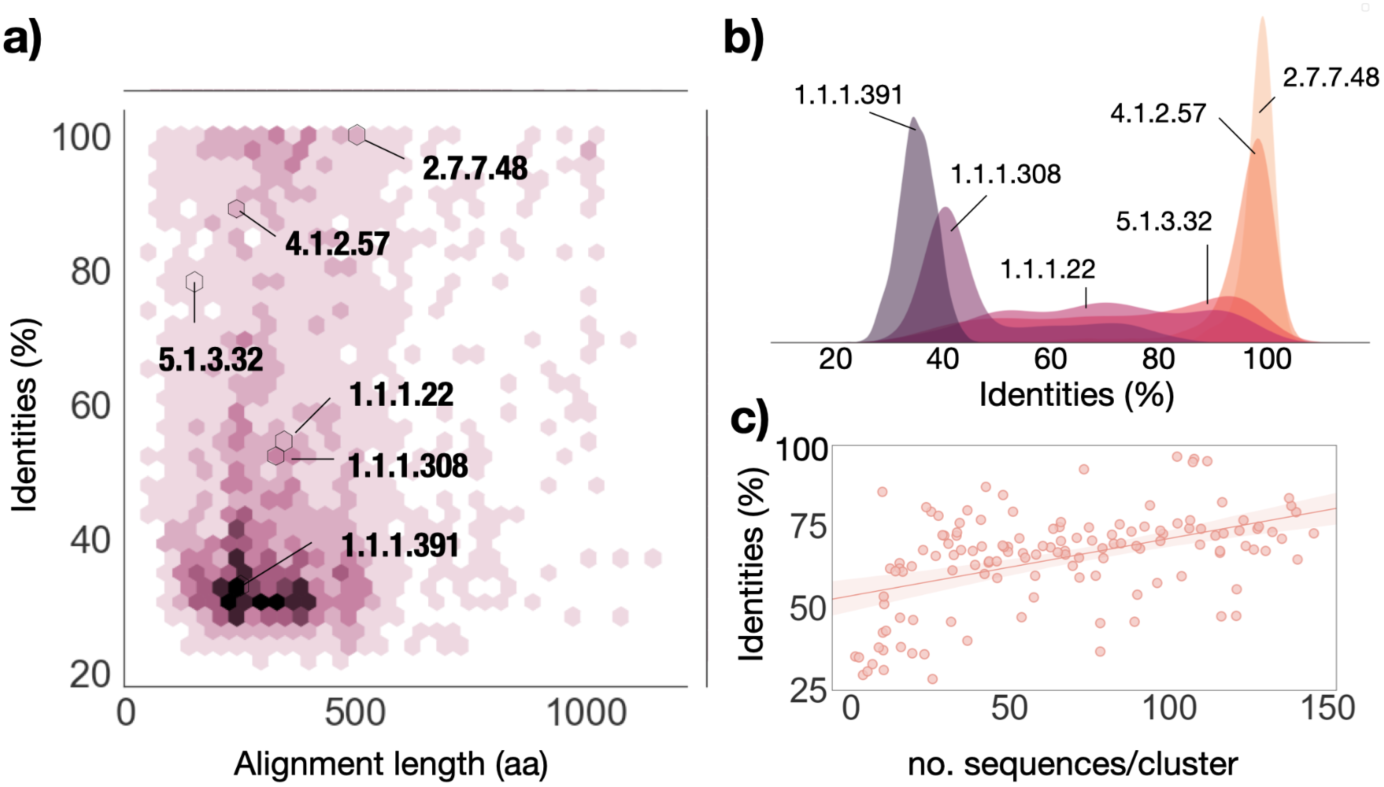
Distance of the generated sequences to the training set. **a)** Identities and lengths for the alignment with the lowest E-value found with MMseqs. Depicted six labels in different regions of the identity plot. **b)** Frequency of generated identities for each of the selected EC classes in a). **c)** Scatter plot between the identities to training set groups for generated sequences and the average number of sequences per cluster for 100 EC classes.

The sequences are distant from the training set, with alignments that show average identities and lengths of 53.1 ± 23.2% and 337.9 ± 151.2 amino acids. BLASTP^35^ searches against the non-redundant protein sequence database provided similar values, with 54.2 ± 22.2% sequence identities (**Fig. S2**). Nevertheless, we observe a non-negligible set of sequences with identities over 90%, with 12.5% of the sequences in the set above that threshold (**Fig. 2a**).

We wondered what are the determinants for these differences in redundancy during generation. Indeed, the model generates sequences distant from the natural space for some classes (1.1.1.391, 1.1.1.308), whereas, for others, they are remarkably close (2.7.7.48) or dispersed (1.1.1.22) (**Fig. 2b**). These sequences span all major enzymatic classes and belong to groups with diverse numbers of members (**Fig. S3**). We thus hypothesised that the differences in these cases may be due to varying degrees of internal redundancy within EC classes. Interestingly, there is a relationship (p-value = 6.04 · 10^−11^, R = 0.54) between the generated sequence identities and the number of clusters per EC class in the training set (**Fig. 2c**), but not so with the total number of sequences per EC class or the total number of clusters (**Fig. S4**). These findings have implications for protein design since specific user cases may require closer (i.e. 90%) or more distant (i.e. <40%) sequences to the natural ones, a property that can be adjusted by fine-tuning the model on specifically clustered datasets.

We then analyzed the predicted globularity, order and functionality of the generated sequences by comparing two crafted datasets of natural and generated sequences (**Methods, Fig. S5**). IUPRED3^36^ revealed that the two datasets show similar globularity levels, with 97.7% and 99.3% of the sequences predicted to be globular (**Table 1**). Average pLDDT values obtained using ESMFold^15^ and Omegafold^37^ also yielded comparable results, with set-averaged pLDDT values of 60 and 85 for the generated and natural datasets, respectively (**Table 1**). To assess the predicted catalytic activity of the generated enzymes, we used HiFi-NN^38^ and CLEAN^39^, state-of-the-art methods for functional annotation from sequence-alone inputs. Both methods accurately annotated a significant proportion of generated sequences, even at the complete EC levels (**Table 1**, 4th EC level), which are more challenging to predict due to the increasing specificity of the described chemical reactions. Specifically, 72% and 34% of the sequences were correctly annotated to the 4th EC level with HiFi-NN, while 56% and 27% achieved correct annotations with CLEAN for the natural and generated datasets, respectively. Given that these generated sequences had not been subjected to prefiltering, the results suggest that the model generates sequences with the potential to catalyse their intended reactions, providing a first solid evidence before proceeding to experimental assessments.

**Table 1:**
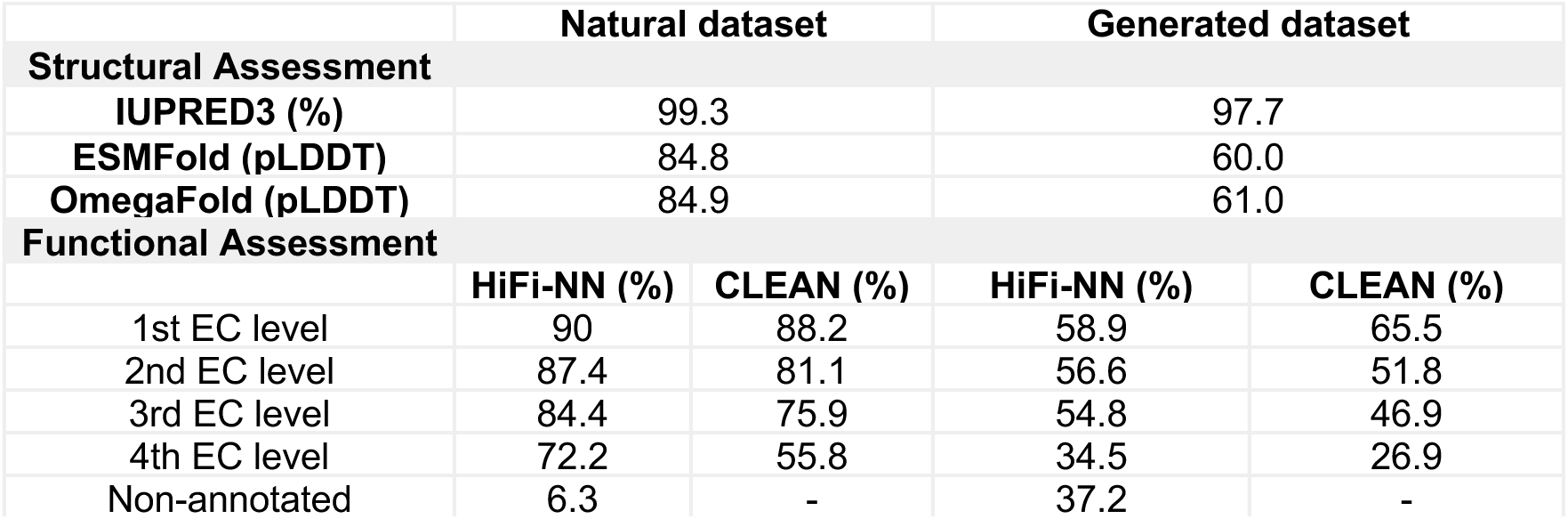
*In silico* structural and functional assessment of ZymCTRL-generated sequences.

## Experimental results

### ZymCTRL generates highly functional carbonic anhydrases

Carbonic anhydrases (EC: 4.2.1.1) represent a group of ubiquitously expressed metalloenzymes that play a crucial role in the rapid conversion of carbon dioxide and water into carbonic acid, protons and bicarbonate ions. From an evolutionary perspective, carbonic anhydrases are remarkably diverse, with eight distinct subtypes (α, β, γ, δ, ζ, η, θ, and ι), exhibiting no similarities in sequence or structure, yet arriving at the same catalytic reaction due to convergent evolution^40^. Kinetically, they have reached catalytic perfection, with a single carbonic anhydrase capable of hydrating one million CO_2_ molecules per second. Beyond their intriguing evolution and chemistry, carbonic anhydrases are attracting significant attention for their potential as biocatalysts in industrial processes. The ever-growing anthropogenic CO_2_ emissions are emphasising the urgency for greener carbon capture and sequestration (CCS) methods to mitigate global warming, with carbonic anhydrases offering a sustainable, cost-effective alternative for CO_2_ fixation^41^.

Given their potential as greener alternatives to CCS processes and being Nature’s fastest enzymes, we decided to focus on carbonic anhydrases, as they confer a stringent yet attractive test for ZymCTRL’s capabilities. To this end, we generated 37,500 sequences with ZymCTRL conditioned on the label ‘4.2.1.1’ (carbonic anhydrases) (**Fig. 3a**). These sequences were generated in zero-shot, i.e., without training or conditioning the model in any way with additional data. We then clustered the sequences to 90% using MMseqs2^42^, ran a BLAST search^35^, selecting those below 60% identity to any hit and filtered by ESMFold’s pLDDT (>70) (**Methods**). The EC class 4.2.1.1 contains representatives of all carbonic anhydrase subtypes, and thus ZymCTRL generates sequences that span from the α to θ classes following that distribution. For this reason, we ran TMalign^43^ against the crystal structure of an alpha (PDB 1CA2) and beta carbonic anhydrase (PDB 1DDZ^44^) (**Fig. 3b**), ensuring we could classify each generated sequence with their subtypes. We then individually inspected all hits to ensure they featured the conserved Zn^2+^-binding sites. Ten beta (CA1-CA5, CA13-CA17) and five alpha (CA6-CA10) carbonic anhydrase were selected for the first round of experimental testing (**Methods, Table S2, Table S5**). Only two of the alpha carbonic anhydrases expressed, but they did it mostly in the insoluble fraction. Conversely, nine of the beta carbonic anhydrases expressed, three were soluble, and one showed mild catalytic activity (**Methods**, **Table S3, and Fig. S6-S7**). While beta carbonic anhydrases are most commonly found in prokaryotes^45^, alpha-carbonic anhydrases are found in mammals. We hypothesised that the observed differences in performance could be due to our choice of the host system (*E. coli*) and the problems that could arise when trying to overexpress sequences that resemble their eukaryotic counterparts in the training set. We therefore focused on designing beta carbonic anhydrases for a second testing round.

**Figure 3:**
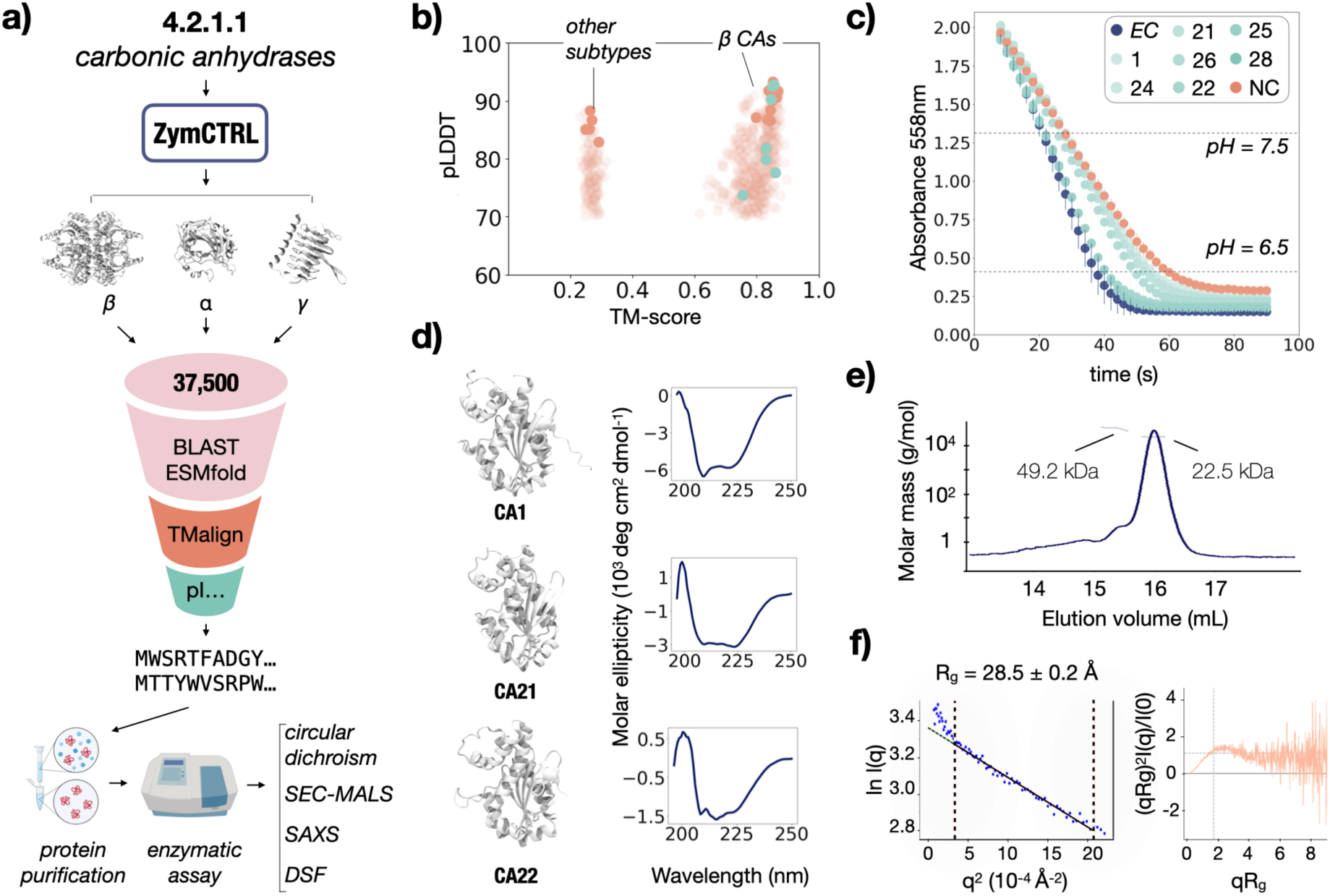
Summary of results for the generated carbonic anhydrases. **a)** Filtering pipeline to obtain the selected hits. Sequences were generated in zero-shot and filtered based on several properties before being assessed for overexpression and purification. **b)** TM align analysis against PDB 1DDZ (beta carbonic anhydrase). The generated enzymes belong to different carbonic anhydrase subtypes. Selected sequences belong to the beta and alpha subtypes. **c)** Spectrophotometric assay measuring the decrease in pH via the indicator phenol red over the course of the reaction. **d)** Circular dichroism spectra and the ESMFold structural predictions for CA1, CA21, and CA22. **e)** Molar mass (g/mol) during SEC-MALS for CA22 and the corresponding absolute molecular weights. **f)** Guinier plot and dimensionless Kratky plot analysis. NC= Negative control; EC = Escherichia Coli.

At the time of our analysis, relevant work was published using different filtering methods^18^. We adopted a similar pipeline and selected ten more beta carbonic anhydrases for a second round of testing. In particular, we ensured that the selected sequences had predicted net charges distant to neutrality and that hydrophobic solvent accessible surface areas were below previously reported thresholds^18^ (**Methods**). In this round, eight of the sequences expressed, and six of them were soluble (**Table 2**, **Fig. 3c, Figs. S8**). We analyzed the activity of these and the previously analyzed sequences with a biochemical assay, additionally including positive and negative controls, namely, the wild-type sequence from *E. coli* (Uniprot ID P0ABE9) and the buffer solution without any target enzyme. In particular, we follow a modified Wilbur-Anderson assay, which measures the time the enzyme requires to decrease the pH of a CO_2_-saturated solution from 7.5 to 6.5 at 0 °C using a colourimetric indicator^46–48^ (**Methods**). Overall, seven of the carbonic anhydrases show activities above a threshold defined by the non-catalysed reaction, with two of them considerably close to the levels obtained by the reference enzyme from *E. coli*. This is a remarkable feat, considering the enzymes are significantly distant from the natural space (non-redundant database, BLAST), with all of them being below 50% sequence identity and the two most active sequences with 39% and 41% sequence identities (**Table 2**).

**Table 2:** Summary of results for the carbonic anhydrase enzymatic assay along with their sequence identity to the non-redundant database and BLAST. Sequences ordered by increasing activity.

| Carbonic Anhydrase | Sequence identity (%) | WAU/mg |
| --- | --- | --- |
| CA1 | 45.97% | 333 ± 15 |
| CA24 | 37.34% | 416 ± 20 |
| CA21 | 42.97% | 437 ± 27 |
| CA26 | 49.28% | 522 ± 58 |
| CA22 | 46.86% | 725 ± 62 |
| CA25 | 41.01% | 1219 ± 28 |
| CA28 | 39.24% | 1310 ± 37 |
| Reference (*Escherichia coli*) | - | 1414 ± 40 |

We further interrogated the enzymes using different biophysical methods. In particular, we analysed CA1, CA21, and CA22 by circular dichroism (CD) and observed that they feature spectra typical of αβ-proteins, as expected from their AlphaFold^49^ predictions (**Fig. 3d**). The analysed sequences appeared to be monomers by size exclusion chromatography (SEC) (**Fig. S9**), and to further confirm these results we subjected most designs to size exclusion chromatography with multi-angle light scattering (SEC-MALS). The results reveal that CA1, CA21, CA22 (**Fig. 3e**), CA25, and CA28 are monomers (**Fig. S10**). Beta carbonic anhydrases exhibit diverse oligomeric states^50^, with the dimer comprising the most common basic unit, although evidence has been presented for monomeric active forms^51,52^. To determine the stability of such monomers, we measured melting temperature (T_m_) for CA22 by Differential Scanning Fluorimetry (DSF). The obtained T_m_ was 47 °C, and the topology of the melting curve agreed with the monomeric state since only one maximum was detected (**Fig. S11**). Due to their higher expressing yields, we further analysed CA1 and CA22 using Small-angle X-ray scattering (SAXS), to further analyse their state in solution (**Fig. S12**, **Fig. 3f**). CA1 and CA22 radius of gyration (R_g_) were 28.23 ± 0.68 Å and 28.45 ± 0.22 Å, respectively, indicating the two proteins exhibit similar sizes. The molecular weight (MW) for the designs was estimated to be within 22.1 - 28.5 kDa, while for CA22, the MW gap was 22.8 - 29.3 kDa, confirming the species behave as monomers in solution. The maximum diameter (D_max_) resulted in 139 Å for both designs. To inspect the putative nature of their folding, we examined the dimensionless Kratky plot (**Fig. 3f**). Such profiles presented the standardised Gaussian bell shape of globular species. Overall, these results confirm that ZymCTRL generates active sequences without the need for additional training in remote regions of the protein space.

### Lactate Dehydrogenases

Lactate dehydrogenases (LDH) (EC: 1.1.1.27) are a well-studied class of enzymes that play a pivotal role in the biotechnology sector, primarily in lactic acid production^53,54^. Their enzymatic role is crucial in the reversible conversion of pyruvate to lactate, with the interconversion of the coenzymes NAD^+^ and NADH^54,55^. The ability of LDH to mediate this conversion underpins the microbial synthesis of lactic acid, an important precursor for the synthesis of chiral compounds such as drugs and pesticides and, more importantly, in the production of bioplastics such as Poly-L-lactic Acid (PLA)^55,56^. PLA is recognised as a biodegradable, eco-friendly alternative to conventional petroleum-based plastics, aligning with the goal of a more circular economy by reducing waste^57,58^. However, its production demands lactic acid of the highest optical and chemical purity. This coupled with the role LDHs play in the food industry, specifically in the development of lactate biosensors for accurate fermentation monitoring and spoilage detection in fermented foods, highlights the importance of efficient enzyme design to enhance lactic acid production^58^.

The ability of ZymCTRL to generate functional enzymes in a zero-shot manner made us wonder to what extent the model would benefit from fine-tuning in a saturated sequence space. Recent work highlighted the uneven sampling of the tree of life in public sequence databases and the impact this can have on the performance of pLMs trained on these datasets^33^. For the purpose of fine-tuning ZymCTRL with sequences sampled from sections of the tree of life absent from public databases, expanding into sequence space beyond the public databases (**Fig. 1b**), we leveraged a proprietary metagenomic graph database (BaseGraph), derived from environmental probes sampled across 5 continents and a 110 °C temperature range.

In particular, we fine-tuned ZymCTRL with a subset of sequences annotated as LDHs from the BaseGraph database, with the majority of these sequences originating from extremophilic organisms extracted from environments with temperatures ranging from −1 °C to 81 °C (**Fig. 4a).** We generated around one million sequences from both the fine-tuned and the original pre-trained model (**Methods**). In line with the notion that language models can benefit from greater protein sequence diversity we observed that, compared to the zero-shot generated sequences, the fine-tuned sequences exhibited higher predicted pLDDT scores and lower sequence identities to the training set (t-test: p < 0.0001 and p < 0.00001, respectively, **Fig. 4b**).

**Figure 4:**
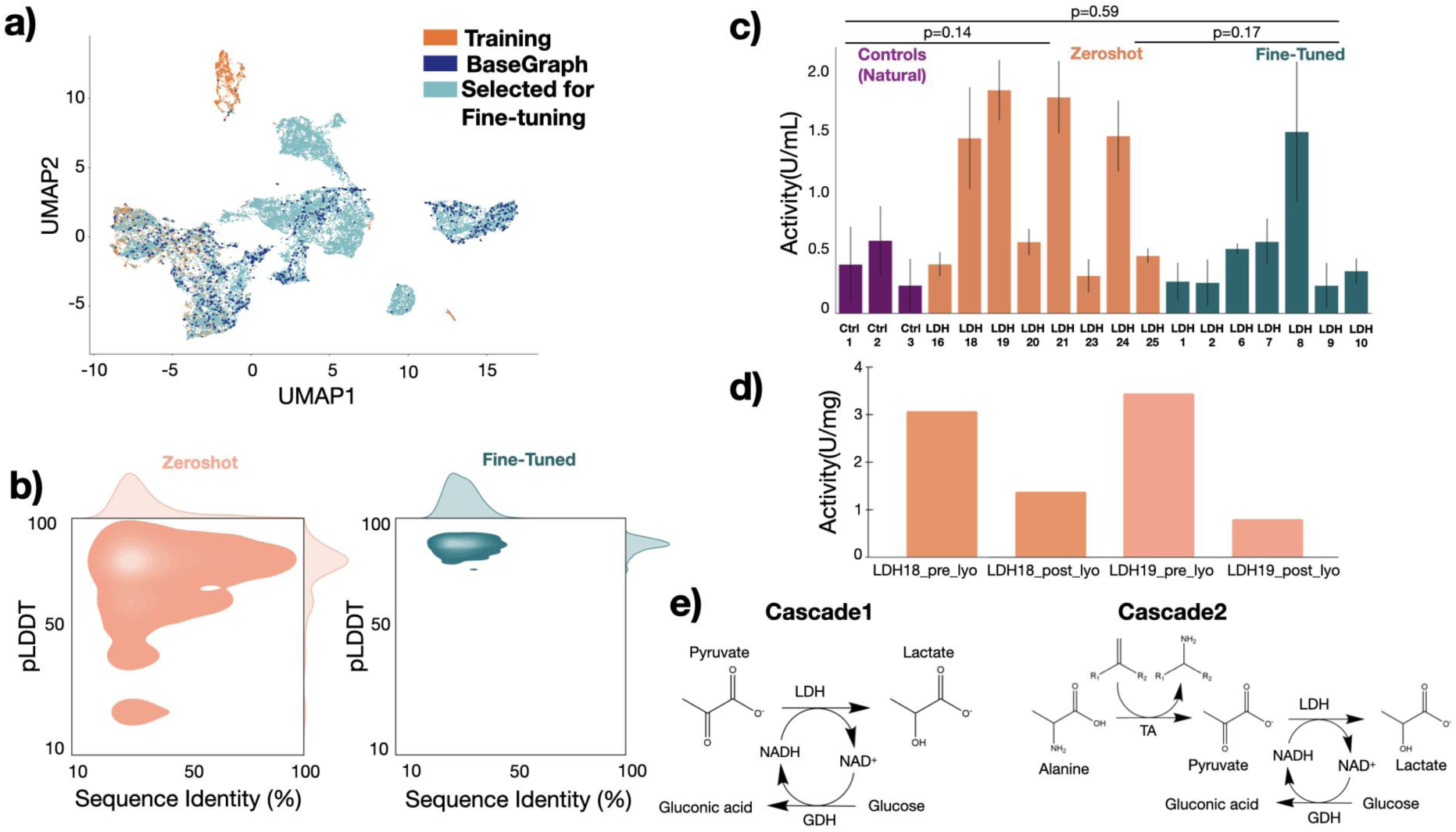
Computational design and functional characterisation of the Lactate dehydrogenases. **(a)** UMAP of ESM-2 embeddings of Lactate dehydrogenases in the training set, BaseGraph and the sequences selected for fine-tuning ZymCRTL **(b)** pLDDT scores against sequence identities (compared to training data) for 756,983 zero-shot (orange) and 1,138,955 fine-tuned (teal) sequences. **(c)** Comparative functional activity of zero-shot and fine-tuned sequences alongside control sequences: Lactate dehydrogenase from Lacticaseibacillus casei and two natural sequences from BaseGraph, used in the fine-tuning process. **(d)** Enzymatic activity of LDH18 and LDH19 pre and post lyophilisation **(e)** One-pot enzymatic cascades, cascade 1: pyruvate is reduced by LDH using NADH as cofactor, while glucose dehydrogenase (GDH) is used to recycle the NADH cofactor forming gluconic acid from glucose. Cascade 2: alanine is employed as the amino donor (coupled to the co-substrate, the ketone acetophenone) for TA and the pyruvate which is generated (together with the co-product (R/S)-α methylbenzylamine, MBA) is reduced by LDH to lactate. Removing the pyruvate serves the dual purpose of driving the reaction and also eliminating pyruvate inhibition of the TA.

**Figure 5:**
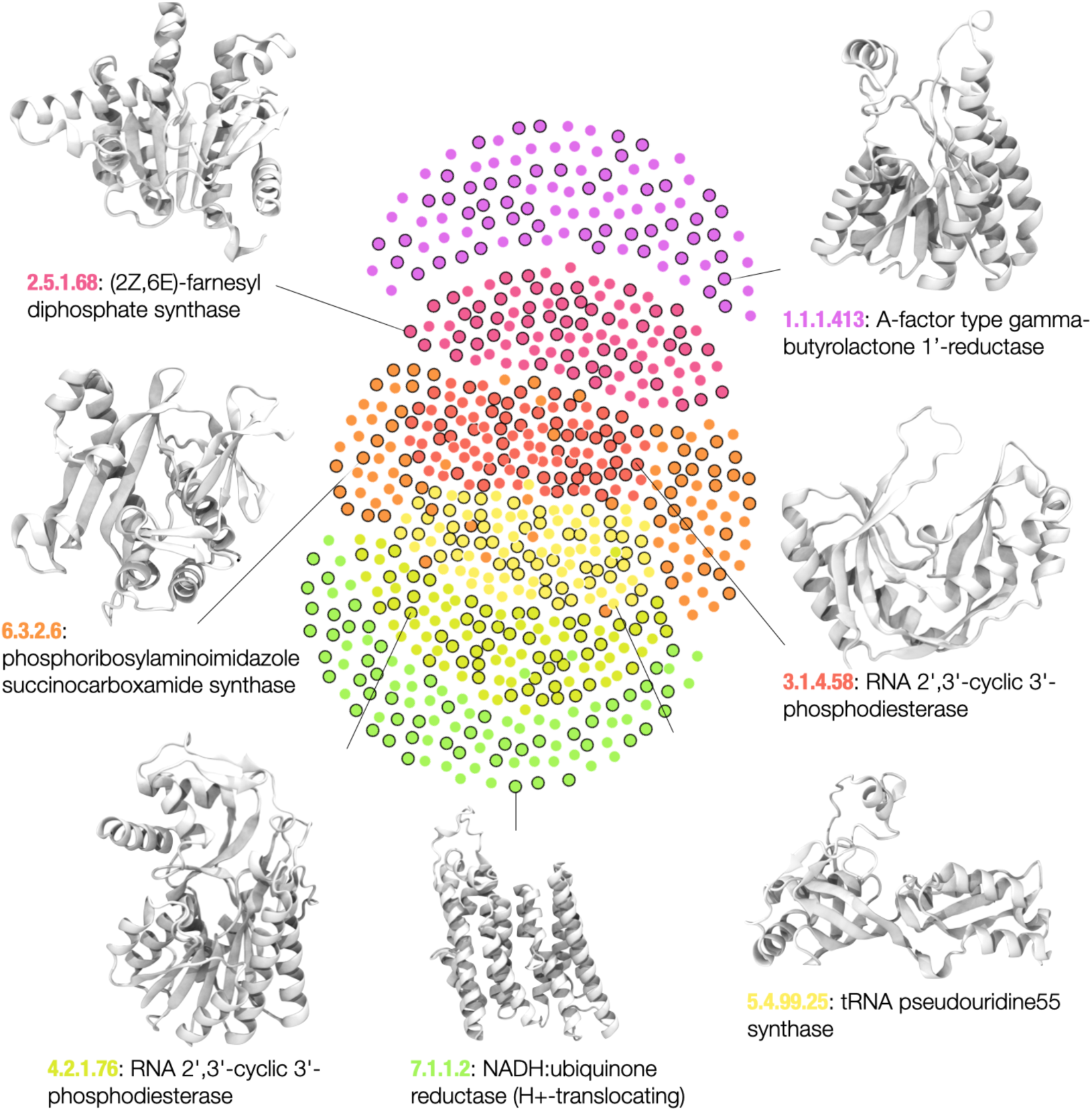
Similarity network of ZymCTRL internal representations defined by cosine similarities. Each node represents a protein sequence either natural or generated (circled in black) from the seven EC classes (EC:1, purple; EC:2: magenta; EC:3, dark orange; EC:4, green-yellow; EC:5, yellow; EC:6, orange; EC:7: green). Nodes are linked whenever their corresponding internal embeddings give a cosine similarity over 0.9.

Given this large pool of fine-tuned and zero-shot generated LDH sequences, filtered and obtained 20 sequences (10 zero-shot and 10 generated) for experimental characterization (**Methods**). Experimentally, all the sequences were expressed in *E. coli* and showed different degrees of solubility (**Fig. S13 - S16 and Table S5, S6, S7**). Notably, fourteen of these sequences displayed measurable enzymatic activity when assayed using a spectrophotometric assay in crude lysates (**Table 3**, **Fig. 4c, Table S4, S5 and S7, Methods**). As positive controls, three wild-type sequences were used, including *L. casei* and two sequences from the BaseGraph database. Negative control consisted of the lysate of an empty pET28a vector (**Methods**). Remarkably, the active sequences demonstrated significant enzymatic activity at a high temperature of 45 °C and across a broad pH range of 4.5 to 9.5 (**Table S7**), offering significant industrial advantages over naturally occurring LDHs, which typically display maximal activity in the narrower, slightly acidic to neutral pH range of 5.5 to 7.0. This exceptional pH tolerance enables robust and versatile biocatalytic processes by allowing the utilisation of a single enzyme across multiple applications with varying pH conditions (**Table S7**). The broad pH optima of these enzymes suggest their robustness in industrial settings, which often encounter pH fluctuations, thereby improving operational stability in large-scale and continuous processes^59–62^. Regarding their activity levels, all three groups (controls, zero-shot, fine-tuned) have comparable performance, with no significant difference observed between either of them (ANOVA f-statistic = 1.99, p = 0.17, n.s., **Fig. 4c**).

**Table 3:** Summary of results for the Lactate dehydrogenase enzymatic assay at pH 4.5. ^a^This value corresponds to pH 9.5 (see Table S7).

| Lactate dehydrogenase | Group | Sequence Identity | Activity (U/mL) |
| --- | --- | --- | --- |
| LDH1 | Fine-tuned | 66.01 | 0.27 ± 0.16 |
| LDH2 | Fine-tuned | 56.06 | 0.26 ± 0.19 |
| LDH6 | Fine-tuned | 65.27 | 0.55 ± 0.05 |
| LDH7 | Fine-tuned | 57.74 | 0.61 ± 0.19 |
| LDH8 | Fine-tuned | 60.70 | 1.56 ± 0.60 |
| LDH9 | Fine-tuned | 58.76 | 0.24 ± 0.19 |
| LDH10 | Fine-tuned | 70.09 | 0.36 ± 0.16 |
| LDH16 | Zero-shot | 74.68 | 0.42 ± 0.10 |
| LDH18 | Zero-shot | 88.32 | 1.50 ± 0.43<sup>a</sup> |
| LDH19 | Zero-shot | No hits found | 1.91 ± 0.25 |
| LDH20 | Zero-shot | 94.04 | 0.61 ± 0.11 |
| LDH21 | Zero-shot | 62.02 | 1.85 ± 0.31 |
| LDH23 | Zero-shot | 96.95 | 0.32 ± 0.14 |
| LDH24 | Zero-shot | 86.23 | 1.52 ± 0.30 |
| LDH25 | Zero-shot | 55.81 | 0.49 ± 0.06 |
| Control1 | *Lactobacillus casei* | - | 0.42 ± 0.30 |
| Control2 | BaseGraph_LDH1 | - | 0.62 ± 0.29 |
| Control3 | BaseGraph_LDH2 | - | 0.24 ± 0.23 |

A critical goal of enzyme engineering is to be able to design materials that can be readily used as biocatalysts in industrial settings. To ascertain whether our designs exhibit such potential, we selected LDH18 and LDH19 for further investigation. We scaled up the production of enzymes LDH18 and LDH19 and proceeded to lyophilisation and integration in enzymatic cascades. Lyophilisation or freeze-drying is a technique for the long preservation of enzymes, bacteria, or other biological products^63^. Main advantages are the preservation of the product and the possibility of rehydration while preserving its activity. While useful, lyophilisation has been shown to often promote molecular changes to the lyophilised proteins, sometimes leading to a complete loss of activity^64^. Interestingly, LDH18 and LDH19 retained activity after lyophilisation (**Fig. 4d**, **Methods**). Following these results, we integrated the enzymes into two different one-pot enzymatic cascades, the first cascade coupled LDH with glucose dehydrogenase (GDH) for cofactor recycling, while the second cascade incorporated a transaminase (TA) to drive the equilibrium toward the desired amine product by removing the pyruvate byproduct mediated by LDH (**Fig. 4e**, **Table S10**, **Methods**). Aiming at testing the enzymes in particularly extreme scenarios, the cascades were assayed at pH 5 and 9.5, with a temperature of 45 °C. Both cascades exhibited mild conversion, with the cascades including LDH18 presenting conversions of 7% and 4% to products, respectively (**Table S8**, **Fig. S17**, **Fig. S18**). While these results indicate that the enzymes require further optimisations, they showcase that enzymes generated without further optimization can be successfully lyophilised and integrated into one-pot enzymatic cascades, displaying initial activities in harsh conditions.

### ZymCTRL embedding space distinguishes enzyme functional classes

We investigated whether the model is capable of discerning the seven EC classes within its internal space. Several techniques are available for reducing the high dimensionality of protein sequences to more manageable, human-understandable dimensions. Recently, manifold learning techniques such as tSNE and UMAP have emerged as powerful dimensionality reduction and visualisation tools^16^. Other studies have focused on hierarchical characterisations^17^, cartesian representations^65^, or similarity networks^18^. Here, we attempted to visualise the model’s embedding space, aiming to understand whether the internal representations of different enzyme classes occupy distinct regions. To this end, we generated ZymCTRL internal representations of natural and generated sequences and constructed a similarity network based on their cosine similarities. The cosine similarity between two vectors—or two enzyme embeddings—measures the similarity between two non-zero vectors defined in an inner product space. Intuitively, two sequences deemed similar by the model will produce larger cosine similarities.

Figure 3 showcases a network containing a total of 700 natural and generated sequences (circled in black), which group together when their cosine similarities surpass the cutoff of 0.9. The enzymes populate well-defined regions, indicating that the model defines sequences belonging to the same class as most similar. Interestingly, there are also inter-class connections, such as oxidoreductases (EC: 1) being linked only to transferases (EC: 2) or translocases (EC: 7) being linked to ligases (EC: 4). These findings reveal that the model has learned an internal space of the enzymatic classes, being able to differentiate the different enzymatic reactions correctly.

## Discussion

The landscape of protein design is undergoing a significant transformation^12,19,20^. While structure prediction tools such as AlphaFold, and numerous studies^22–24,66^ have demonstrated the ability to design tailored protein structures with remarkable accuracy, the next frontier lies in designing proteins with customized functions. Recent years have seen AI methods emerge as powerful tools for enzyme design, with already several successes in creating enzymes with catalytic efficiencies^22–24,66^. Inspired by the versatility of language models in fulfilling user-defined requests, we trained ZymCTRL, a protein language model tailored for designing enzymes that catalyze user-specified reactions. ZymCTRL can generate high-quality enzymes even for reactions with limited known examples without further training, a property that will allow the augmentation of isolated enzyme families in a cost-effective fashion. Additionally, fine-tuning on curated datasets allows for control over the properties of the generated sequences.

We tested ZymCTRL’s robustness for two different enzyme families. Firstly, we generated carbonic anhydrases in zero-shot, without additional conditioning of the pre-trained model with external data. Of the 20 generated beta carbonic anhydrases, seven displayed activity following purification, with two close to the wild-type positive control. Biophysical analyses confirmed well-folded, globular designs. These results are particularly noteworthy considering the challenging nature of carbonic anhydrases as a test for the model, the sequences’ significant divergence from natural sequences (<45%), and the fact that the sequences were generated in zero-shot, with a filtering pipeline executable on a standard GPU within hours. Furthermore, we also generated lactate dehydrogenases in zero-shot, attractive enzymes for lactic acid production. Many of the generated sequences surpassed the controls, and two were successfully scaled up, lyophilised, and integrated into enzymatic cascades, retaining activity and exhibiting conversion.

Motivated by these promising results and recognizing the imbalance in training data across labels and the tree of life, we fine-tuned the model on a diverse set of lactate dehydrogenases from BaseGraph. Compared to their zero-shot counterparts, we observed that post-fine-tuning, generated sequences were more likely to explore novel sequence space and exhibited higher average pLDDT scores. These findings further demonstrate the potential of enhancing the model’s knowledge with user-curated datasets, thereby enabling control over various properties such as distance to the training set (Fig. 2c) and other attributes like thermostability.

Despite the success demonstrated in experimental settings, there are potential challenges to address in the future. Here, we have shown that conditional generation works for enzyme classes with narrow substrate scopes. However, certain EC classes encompass multiple substrates (such as hexokinases, EC: 2.7.1.1), posing a current challenge in generating sequences tailored to individual substrates within these classes. Future efforts will involve encoding substrate information during fine-tuning of sequences for individual substrates or training models capable of generating sequences with control at the substrate level. Moreover, an aspect that requires further investigation is the variability in solubility and activity among the sequences. For example, the lactate dehydrogenases show similar results regardless of whether they were subjected to a filtering process or randomly generated (**Table 3, Methods**). Understanding these differences, possibly by using explainable artificial intelligence techniques, will be crucial for further improving the efficiency of these models and expanding our limited understanding of sequence-function relationships.

While there are a few challenges ahead, our findings highlight the potential of conditional language models as potent tools for the controllable design of enzymes, paving the way for a myriad of applications. To the benefit of the community, we make code, model weights and training data publicly available.

## Acknowledgements

The authors gratefully acknowledge the scientific support and HPC resources provided by the Erlangen National High-Performance Computing Center (NHR@FAU) of the Friedrich-Alexander-Universität Erlangen-Nürnberg (FAU) under the NHR project b114cb (UID 210235). Federal and Bavarian state authorities provide NHR funding. NHR@FAU hardware is partially funded by the German Research Foundation (DFG) – 440719683. N.F. acknowledges support from an AGAUR Beatriu de Pinós MSCA-COFUND Fellowship (project 2020-BP-00130) and a Ramón y Cajal contract RYC2021-034367-I. U.E acknowledge support from a Ramón y Cajal contract RYC2020-029773-I. We thank the generous funding from the L’Oreal-UNESCO For Women in Science Price 2022. We thank Florian Grün for stimulating discussions. We thank Ahir Pushpanath, Neem Patel, Marcus Leung, Sergio Romero-Romero and Marc Garcia-Borràs for their helpful feedback and inspiring discussions. We are grateful to Mònica Buxaderas at the Automated Crystallisation Platform (PAC) of the Scientific Parc of Barcelona during SEC-MALS experiments. We acknowledge support from Carlo Carolis and Miriam Alloza at the Protein Technology Unit of the Centre for Genomic Regulation (Barcelona, Spain). The authors gratefully acknowledge the biodiversity stakeholders around the world who have granted Basecamp Research permissions to access their sites for the collection and analysis of environmental samples for metagenomic research and commercialisation purposes.

## Contributions

N.F and S.L trained the model. G.M and N.F fine-tuned and filtered sequences for the different testing datasets. S.F tested the LDH designs. R.I-V performed the scaled-up and biophysical assays for the carbonic anhydrases, with the valuable input from U.E. I.T.N, L.S.S and S.S performed the carbonic anhydrases activity assay. G.A performed the HiFi-NN calculations. N.F and G.M wrote the manuscript with the aid of P.L, S.F, U.E and R.I-V. N.F and P.L designed the work and coordinated the teams.

## Methods

### Dataset preparation and vocabulary encoding

We downloaded sequences in the Uniprot database which had an EC number annotation through the UniProt web interface (July 2022, version 2022.1) giving a total of 37,624,812 sequences. To avoid multi-chain sequences with multiple EC assignments, we removed sequences with several EC labels, giving a total of 36,276,604 sequences. We split the database into training (90%) and evaluation (10%) datasets. We used a block size of 1024, separated control tags and sequences with a separator token, and further specified the boundaries of sequences below 1024 amino acids with start and end tokens. We fit as many complete sequences as possible in the 1024 window, provided that sequences are not split across blocks. The sequences will follow this schema if their length fits in the 1024 window: <control tag><sep><start><ENZYME SEQUENCE><end><|endoftext|>, and the following scheme otherwise: <control tag><sep=<ENZYME SEQUENCE=<|endoftext|>. Sequences over 1024 amino acids (∼3%) were truncated to the N-terminal part.

### Vocabulary encoding

We train our model with an associated label (control tag) per sequence. Following recent studies^7,21^, we tokenised our enzyme sequences using amino acid encoding. We further tokenised the labels in the dataset, to account for similarities among sub-classes in the same classes and help the model generalise in lower-populated catalytic reactions. This way, the control tag ‘1.1.1.1’ is split into its categories (‘1’ + ‘.’ + ‘1’ + ‘.’ + ‘1’ + ‘.’ + ‘1’) and shares 6 tokens with ‘1.1.1.2’.

### Model pre-training

We use a Transformer decoder model as architecture for our training. The model uses the original dot-scale self-attention^22^. The architecture uses that of the CTRL/GPT2 Transformer, which was downloaded from HuggingFace^23^. ZymCTRL consists of 36 layers, a model dimensionality of 1260, and 16 attention heads. The model was optimised using Adam (β1 = 0.9, β2 = 0.999) with a learning rate of 0.8e-04, following previous works^24^. A batch size of 4 per device was used accumulating 4 gradient steps, resulting in a total batch size of 768. We trained for 179,000 steps on 48 NVIDIA A100s 80GB in 15,000 GPU hours. Parallelism of the model was handled with DeepSpeed^25^.

### Perplexity evaluation

We evaluate the perplexity as the exponentiated average negative log-likelihood of a sequence. Because ZymCTRL is a fixed-width causal language model, we evaluate the perplexity using a previously described sliding-window strategy (https://huggingface.co/docs/transformers/en/perplexity). In particular, for a tokenised sequence (x = x_0_, x_1_, x_2_, x_3_… x_n_), the perplexity of x is evaluated as:

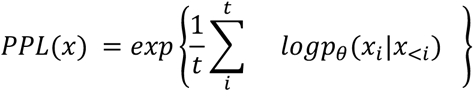

where *logp_θ_*(*x_i_*|*x*_<*i*_) is the log-likelihood of the i-th token conditioned on the *x*_<*i*_ token.

### Dataset creation

We randomly sampled two sequences per EC number for multi-sequence classes and one sequence otherwise. For the generated dataset, we then generated 20 sequences per EC number and selected the best or two best perplexity-scoring sequences depending on the number of sequences in the equivalent natural dataset’s class. Each dataset contained 11,438 sequences. We ensured that the generated sequences followed the natural dataset length distribution (**Fig. S2**), by applying a length limit of 600 to all labels, except when no sequence could be generated at that length, hence the limit was extended to 1024. In all cases, the sequences were only selected if they had been finished and not truncated by the model by ensuring the generation of the end-of-sentence token.

### Functional prediction analyses

#### CLEAN

We ran CLEAN^39^ on the generated and natural datasets, each comprising 11,438 sequences, using the maximum separation inference method. This method, chosen for its consistent precision and recall performance, employs a greedy algorithm that preferentially selects EC numbers exhibiting the greatest pairwise distance from other EC numbers relative to the query sequence^39^. To assess label accuracy, we compared the predictions made by CLEAN against the labels of the generated and natural sequences across the EC hierarchy.

#### HiFi-NN

We annotate both the generated and natural datasets using HiFi-NN^38^ with the set of sequences in Swissprot which have an EC as the lookup set. By default, HiFi-NN assigns annotations to every sequence in the set. To remove potentially spurious annotations, we use a cosine distance (1 - cosine similarity) cutoff of 0.3^38^. As before, we check if the desired EC exists in the set of annotations assigned by HiFi-NN. This is performed across each level of the EC hierarchy.

### Amino acid propensities

We computed the natural amino acid propensities by taking sequences from the ten most populated EC classes in the training dataset. We generated sequences from the same EC numbers (**Fig. 1b**), with 20 sequences per parameter set. We tested a sampling generative procedure^26^, with a temperature of 1, max_length of 1024, top_p of 1, repetition penalty 1.2 and 1.3, and top_k for the values from 5 to 20, and 30, 50, 100, 200, and 458. Accuracy to match the natural distribution was computed as the sum of the absolute differences between all amino acid pairs. Repetition penalty of 1.2 provided better results in all cases. Top_k=9 gave the closest distribution to the target propensities (**Fig. S1**).

### Model finetuning

Additional metagenomic sequences for fine-tuning were derived from Basecamp Research’s graph database. Environmental samples subjected to metagenomic sequencing & chemical analysis were collected after receiving landowner’s permission and entering access-benefit-sharing agreements with the relevant local or national authority, following Nagoya protocol guidelines. All samples were sequenced with both long-read (Oxford Nanopore GridION) and short-read (Illumina NovaSeq 6000) sequencing methods applied to each sample after extraction. Following standard sequencing QC, an assembly-based approach was followed, generating *de novo* assemblies that were subjected to polishing and open-reading frame annotation. All open reading frames were annotated functionally *in silico* with a custom annotation pipeline. Translated protein sequences alongside functional, genomic and sample information were inserted into Basecamp Research’s graph database. For fine-tuning, we first extracted all sequences from the graph database that were annotated with ‘EC:1.1.1.27’ (lactate dehydrogenases). We then clustered these sequences at a 50% identity threshold using the MMSeqs2 algorithm applying a length filter of 240-380 amino acids. A subset of 973 sequences were randomly selected for fine-tuning. For fine-tuning, we started from the pre-trained ZymCTRL model, adjusted the learning rate to 0.8e-06 and continued training the model for an additional 100 epochs, at which point the training had plateaued. The final model weights obtained at the end of this fine-tuning process were saved and used for inference.

### Selection of carbonic anhydrases

2000 generations were performed each producing 20 sequences, which is the maximum number of sequences that fit into a single NVIDIA A40 per generation call. Sequences that did not produce an end-of-sentence token were discarded, along with duplicates. This produced 37,503 sequences. BLAST searches and subsequent filtering at <60% yielded 1,464 sequences of interest. We further filtered those sequences based on ESMFold pLDDT values (>70), producing 479 sequences and then ranked the sequences using TM-align against the alpha and beta carbonic anhydrases template PDBs 1CA2 and 1DDZ, respectively. For the first ranking we selected five alpha and ten beta carbonic anhydrases following the order of the ranking and ensuring the designs had the necessary catalytic pockets and the correct ProteInfer^67^ and Interproscan^68^ predictions. For the second filtering process, we added 130 extra sequences with higher BLAST identities to the original 479 sequences. From this set, we discarded sequences with net charges in the interval (−2,+2) and whose hydrophobic solvent accessible surface area calculation yielded values higher than those expected from an idealised monomer. In particular, we followed previous work^18^ and discarded sequences whose hydrophobic SASA was 1.7 times the “ideal surface” computed using the ideal sphere for the same length protein. The ideal surface was computed as in Dill *et al.*^69^, by defining the idealised radius of a protein based on its number of residues as 2.24*(no. residues^0.392). We computed pI values using biopython^70^ and removed variants which had values too close to a pH of 7. Sequences were further manually inspected to ensure they contained the catalytic residues. Ten extra sequences proceeded for experimental testing.

### Expression of carbonic anhydrases

Carbonic anhydrases sequences cloned in pET-28a(+) downstream the six-histidine tag were expressed in BL21(DE3) *E. coli* strain, previously transformed with the pGro7 chaperone-encoding plasmid from the TaKaRa Chaperone Plasmid Set (Cat. #3340, Takara Bio Inc.). Cells were grown in LB medium supplemented with kanamycin sulphate (50 µg/mL), chloramphenicol (34 µg/mL) and L-arabinose (500 µg/mL) at 37 °C and 200 rpm of agitation until the optical density at 600 nm (OD_600_) reached 0.6. Isopropyl β-D-thiogalactoside (IPTG) and ZnSO_4_ were added at a final concentration of 0.1 mM and 0.5 mM, respectively, and protein expression was carried out at 30 °C for four hours. Finally, cells were harvested by centrifugation at 4000 g for 30 minutes at 4 °C and flash-frozen in liquid nitrogen for storage at −80 °C. Such an expression protocol was established after a thorough screening of protein production conditions. In particular, various growth temperatures (30 °C, 37 °C), OD_600_ before induction (0.5, 0.6), IPTG concentrations (0.1 mM, 1 mM), and incubation temperatures and times (30 °C for 4h, 23 °C for 4h, 18 °C for 4h, 16 °C overnight, 13 °C overnight, 13 °C for 4h, 13°C for 4h) were screened. Cell pellets were resuspended in one fifth of the initial culture volume using 20 mM Tris equilibration buffer (containing 20 mM Tris, 150 mM sodium chloride, and 20 mM imidazole, pH 8.8) and subjected to sonication using a QSonica 4-probe horn. Sonication parameters were set to 5 minutes at 50% amplitude with alternating cycles of 10 seconds on and 10 seconds off. The lysates were then clarified by centrifugation at 3900 g for 30 minutes at 4 °C. Subsequently, the soluble fractions were carefully transferred to 15 mL centrifuge tubes, while the no pellets were retained for analysis of the target protein in the insoluble fractions. The successfulness of the protein overexpression trials was checked in mPAGE® 12% Bis-Tris Precast Gels.

### Purification of carbonic anhydrases

To assess the activity of carbonic anhydrases, a small-scale production and purification were carried out. For protein purification, a His60 Ni Gravity disposable polypropylene column loaded with 1 mL HisPur™ Ni-NTA Resin (ThermoScientific^TM^) and all buffers were equilibrated to the working temperature. The column was washed with 10 column volumes of Equilibration Buffer (20 mM Tris, 150 mM sodium chloride, 20 mM imidazole, pH 8.8), then 2 mL of clarified lysate was added. The column was sealed and gently inverted for 1 hour at 4°C to allow binding of the target protein. After repositioning the column vertically to settle the resin, a stand with clean empty tubes was placed under the outlet for fraction collection. The flow was initiated by removing the stoppers and collecting 1 mL fractions. The column was washed with 10 column volumes each of Equilibration and Wash Buffer (20 mM Tris, 150 mM sodium chloride, 40 mM imidazole pH 8.8), followed by elution of the target protein with 10 column volumes of Elution Buffer (20 mM Tris, 150 mM sodium chloride, 300 mM imidazole, pH 8.8), with fractions collected and analysed by SDS-PAGE and Bradford protein assay to monitor protein concentration. Excess imidazole was removed from the required fractions using a PD-10 desalting column packed with Sephadex G-25 resin (Cytiva^®^) for downstream applications and stored in Storage buffer (20 mM Tris and 150 mM sodium chloride at pH 8.8).

For biochemical and structural characterisation, a high-scale production and purification took place. Cell pellet from a 2 L culture was resuspended in 80 mL of lysis buffer (50 mM TRIS-HCl pH 8, 150 mM NaCl, 1 mM DTT, 20 mM imidazole, 1X cOmplete™ EDTA-free protease inhibitor cocktail, 20 µg/mL DNAse) and sonicated on ice with a Branson 250 Digital Sonifier TM (MarshallScientific) bearing a 1.5 mm tip at 25% amplitude in 10 seconds pulse alternated with gaps of 10 seconds for a total of 8 minutes. Then, complete lysis was achieved in a CF-1 cell disruptor (Constant System Ltd.) at a pressure of 1.36 kbar at 4 °C. The protein suspension was clarified by centrifugation at 48000 g for 1 hour at 4 °C and the supernatant filtered with a 0.22 µm pore limit. The purification of carbonic anhydrases CA22, CA25 and CA28 was performed in an ÄKTA pure™ 25 (Cytiva^®^) following a standardised protocol which consisted of a first immobilised metal affinity chromatography (IMAC) in a HisTrap HP 5 mL (Cytiva^®^) by an imidazole gradient in 20 column volumes (CV), an anion exchange chromatography (AEx) in a Capto HiResQ 5 50 (Cytiva^®^) by a NaCl gradient in 40 CV, and a final size-exclusion chromatography (SEC) using a Superdex 200 Increase 10/300 GL (Cytiva^®^). In the case of the IMAC, buffers A and B had 50 mM TRIS-HCl pH 8, 150 mM NaCl, 1 mM DTT, and imidazole, whose concentration was 20 mM and 400 mM for buffers A and B, respectively. AEx buffers C and D harboured 20 mM TRIS-HCl pH 8, 2 mM DTT; buffer C had no NaCl whereas buffer D presented 1 M NaCl. The SEC buffer solution was 10 mM TRIS-HCl pH 8, 500 mM NaCl, 2 mM DTT. Carbonic anhydrase CA1 was subjected to a similar protocol, but excluding the AEx step and switching TRIS pH 8 by HEPES pH 7 since its theoretical pI is 8.3. To ensure carbonic anhydrases had been isolated, gel bands were cut and analysed by nLC-MS/MS at the Service of Genomics and Proteomics from the Center for Biological Research (CIB-CSIC, Madrid, Spain). Once pure, the selected fractions of carbonic anhydrases were concentrated in a Vivaspin® Turbo 10 MWCO (Sartorius), 10% glycerol was added, and proteins were flash-frozen in liquid nitrogen for storage at −80 °C.

Purification of carbonic anhydrases CA21, CA24 and CA26 took place at the Protein Technologies Unit from the Centre for Genomic Regulation in Barcelona, Spain. Once the cell lysate was centrifuged and the supernatant filtered, for each protein a HisTrap HP 5 mL (Cytiva®) was loaded. Then, the column was washed with 15 CV at 100% A, and elution took place in two steps at 10% B (10 CV) and 100% B (5 CV). Fractions with the highest CA concentration were directly used for SEC-MALS characterization in the same facility.

### Circular dichroism

CD results were acquired from 150 μL of sample volume in a 1 mm quartz cuvette with a Jasco-815 (Jasco Analitica Spain) spectropolarimeter from the Scientific and Technological Centers of the University of Barcelona (CCiTUB). Several sample concentrations were tested, including 1, 3, and 6 mg/mL; and the buffer of choice was 1X PBS pH 8, 1mM TCEP. The blank sample was taken from the ultrafiltrate of the protein concentrator. The selected wavelength range was 190-300 nm in 1 nm bandwidth. Spectrum Measurements software was used for data acquisition, Spectra Analysis software for data processing and BestSel^71^ online server for estimation of secondary structure content. Figures represent mean residue molar ellipticity after buffer subtraction ([Θ]), with [Θ] = Θ/l·c·Nr, where Θ is the ellipticity signal in millidegrees, is the cell path in mm (1mm), c the molar protein concentration, and Nris the number of amino acids per protein including histidine tags and thrombin cleavage site^72^.

### Melting temperature

Differential Scanning Fluorimetry (DSF) was performed in a iCycler iQTM Real-Time PCR Detection System (Bio-Rad) with opaque white plates of 96-conical wells and a transparent film cover, using SYPRO^TM^ Orange as a fluorophore at a final concentration of 5X in a total volume of 25 ρL. Before setting the reaction, the plate was preheated at 95 °C for 30 minutes to minimise background signal. The DSF protocol started equilibrating the plate with the samples at 20 °C for 5 minutes, then temperature was increased by 1 °C, and the plate was incubated for 45 seconds at each temperature until 95 °C were reached, with a final incubation of 5 minutes. For acquisition, the excitation wavelength chosen was 495 nm and emission was recorded at 519 nm.

### SEC-MALS

SEC-MALS measurements for CA1, CA22, CA25 and CA28 were performed at the Automated Crystallisation Platform (PAC) from the Scientific Parc of Barcelona (Spain) in a Superdex 200 Increase 10/300 GL column (Cytiva®) mounted on a FPLC system (Shimadzu Prominence) coupled to a scattering DAWN-HELEOS-II-detector (Wyatt Technology®), followed by a Optilab T-rEX dRI (Wyatt Technology ®) refractometer. The injected sample volume for each carbonic anhydrase was 100 μL, and the running buffer of choice was the same as for the SEC purification step. Data treatment and the corresponding calculations were done within the ASTRA software.

### Carbonic anhydrase enzymatic assay

A spectrophotometric assay was developed to monitor consumption of CO_2_ using the following downstream enzyme cascade (modified Wilbur-Anderson assay^73^). During CO_2_ hydration, protons are released causing a decrease in the pH of the solution which is measured using a pH indicator (phenol red). In this assay, 1 μM for each target protein was incubated with 100 μl reaction buffer (20 mM TRIS containing 200 μM phenol red and 1 mM ZnSO_4_, pH 8.8) on ice. Subsequently, 100 μl of CO_2_-saturated water were added into the solution, and transferred quickly into a well of a 96 well plate. Change in absorbance was monitored at 558 nm and recorded every 2 seconds from second 8 (CO_2_ mixing time) using BioTek Epoch^TM^ microplate spectrophotometer. As the pH change was indirectly measured via the absorbance change, the absorbance values corresponding to pH 7.5 and pH 6.5 were required for the calculation. These values were predetermined by measuring the absorbance of a mixture of 100 μL of reaction buffer and 120 μL of deionised water with the pH adjusted to 7.5 or 6.5 using HCl. The absorbance values corresponding to pH 7.5 and pH 6.5 were 1.3 and 0.4, respectively (**Fig. 4d**). Enzyme activity is expressed in Wilbur-Anderson units (WAU), i.e., one WAU measures the pH to drop from 7.5 to 6.5 at 0 °C. The obtained times for the enzyme sample and blank were designated t and t_0_, respectively. In our experiments, the t_0_ value was determined to be 30.5 s when TRIS buffer was used as the blank buffer. The Wilbur Anderson units (WAU) were calculated as follows:

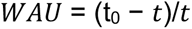

Specific activity was expressed as WAU/mg of enzyme.

### Western blots

Proteins resolved via SDS-PAGE were electro-transferred to nitrocellulose membranes using a Bio-Rad Trans-Blot SD semi-dry apparatus, set at 11 V and 200 mA for 35 minutes, using Towbin buffer as the transferring medium. Post-transfer, membranes were blocked in PBST-milk for 1 hour with gentle agitation, then probed with 30 mL of PBST-milk containing 3 µL of anti-His-Tag primary antibody (His-Tag Mouse anti-Tag, 1B7G5, Proteintech) for 1 hour. Following primary antibody binding, membranes were washed three times with PBST-milk for 5 minutes each. Subsequently, membranes were incubated with 30 mL of PBST-milk supplemented with 6 µL of secondary antibody (Goat Anti-Mouse IgG H&L (HRP), ab205719, Abcam) for 1 hour, followed by three 5-minute washes in PBS. Detection was performed using a substrate mixture consisting of 1 mL Bio-Rad Opti-4CM diluent, 0.2 mL Bio-Rad Opti-4CM substrate, and 9 mL deionised water, with a 20-minute incubation to allow band development.

### SAXS

Small Angle X-ray Scattering (SAXS) data for CA1 and CA22 were obtained at the beamline BM29 from the European Synchrotron Radiation Facility (ESRF) in Grenoble, France. SAXS profiles were processed with BioXTAS RAW 2.2.2 software, and size and shape parameters were calculated with the ATSAS 3.2.1 package.

### Selection of Lactate dehydrogenases

For the sequence selection of the fine-tuned sequences for testing, we applied a length filter of 240-380 amino acids, aligning with the length distribution of naturally occurring LDHs, to reduce the number of sequences for selection. Subsequently, we used MMSeqs2 to cluster the sequences at 90% similarity and 80% coverage. The sequences were further filtered by applying a perplexity threshold of 1.5, followed by re-clustering at a 50% identity threshold using MMSeqs2, to further reduce the number of sequences for selection resulting in a total of 385,817 sequences. For the final selection of LDH sequences to be experimentally tested, the same criteria that was used for the carbonic anhydrases was applied. Thresholds for net charge, solvent-accessible surface area, and isoelectric point were determined from the distribution observed in 100 natural LDH sequences. In addition to passing these physicochemical property filters, sequences were further filtered based on log likelihood scores from the ProteinMPNN model, selecting sequences that had to meet a minimum log likelihood score threshold of 1.2 to 2.0. Additional structural criteria filters included the use ESMFold’s pLDDT (>70) and a TMalign score (>0.85) when compared to the crystal structure of the thermophilic *Bacillus stearothermophilus* LDH (PDB:1LDB). The final selection of sequences were randomly chosen from the top 100 sequences that passed all the filters. For zero-shot generation, we used ZymCTRL to generate 756,983 sequences. Two of the sequences were subjected to the same filters as the fine-tuned subset (LDH17 and LDH20), while another seven were selected randomly from the entire set. This gave a total of 10 sequences.

### Expression of Lactate dehydrogenases

LDH sequences cloned in pET-28a(+) were expressed in BL21 (DE3) *E. coli* strain. Expression tests were performed at three temperatures: 20°C, 25°C and 30°C, to determine the optimal conditions for expression. Expression was carried out in 2 mL cultures in 48 deep well plates using TB medium supplemented with 0.1 M potassium phosphate buffer (KPi) and kanamycin sulphate (50 µg/mL). After 4.5 hours of growth at 37 °C with 700 rpm shaking (OD_600_ = 0.6-1), expression was induced by adding 0.5 mM IPTG and incubated for further 18 hours at the selected temperatures. For expression analysis, 100 μL of the induced culture was subjected to sonication in a Polyethylene Glycol (PEG) bath (Q Sonica) set to 50% amplitude with a 2 minute 30 second total run time (10 sec on, 10 sec off pulsing). Both total and soluble protein fractions were analysed by SDS-PAGE, loading 8 μL of a 15 μL aliquot of each fraction onto the gel. Once the best expression temperature was determined, the remaining cultures were centrifuged at 4000 rpm for 20 min at 4°C, the pellets of the corresponding best expression temperature were resuspended in 0.25 mL of lysis buffer (0.1 M KPi pH 7), sonicated twice as previously described in a PEG bath and the samples were centrifuged at 10000 rpm for 10 min and supernatant (∼200 µL) collected for further analysis. LDH18 and LDH19 were expressed at larger scale following a similar procedure. 200 mL of TB medium supplemented with 0.1 M KPi and kanamycin sulphate (50 µg/mL) were inoculated with 2 mL of pre-culture. When OD_600_ = 0.6-1 was reached, the expression was induced with 0.5 mM IPTG and the flasks were moved to 25°C for 18 hours. The cells were centrifuged at 4000 rpm for 20 min at 4°C, resuspended in a lysis buffer and sonicated with a CV18 probe for 4 min at 60% amplitude (5 sec on, 2 sec off pulsing). The samples were centrifuged for 30 min at 10000 rpm at 4°C and the supernatant was frozen at −80°C for 3 hours before lyophilisation, performed with an Advantage Pro freeze dryer (SP Scientific).. 50 µL of lysate was stored as liquid to compare the activity before and after lyophilisation. SDS-PAGE was performed as described for small scale expression samples.

### Lactate dehydrogenase enzymatic assay

One unit of activity is defined as the amount of enzyme which catalyses the conversion of 1 µmol of substrate per minute under standard conditions. The LDH activity was tested at 25 °C in 200 µL of 0.1 M KPi pH 7, 0.15 mM NADH, 10 mM pyruvate. The reaction was initiated by adding 5 µL of LDH lysate or 5 µL of an empty pET28a lysate for negative controls. The assay was performed in the same manner at different temperatures (25 and 45 °C), pHs (4.5 and 9.5), cofactor (NADPH) and substrate concentration (50 mM). The consumption of NADH was followed spectrophotometrically at 340 nm (NADH ε_340nm_= 6220 M^1^ cm^1^). 96-well plates and a microplate spectrophotometer (Multiskan GO, ThermoScientific^TM^) were used to detect the reduction of the signal at 340 nm. For thermal denaturation, LDHs were incubated at 60 °C for 10 min, cooled on ice for 5 min, and then residual enzymatic activity toward pyruvate was assayed as described above. To obtain the final activity values for each enzyme, the background activity from the pET28a vector was subtracted. This approach ensures that the reported enzyme activities reflect the true contributions of the enzymes of interest, rather than any inherent background signal or activity present in the expression system. All activities were performed at least in triplicate and reported values represent means ± standard deviation (s.d.).

### Experimental protocol for cascade 1

The first cascade reactions were conducted in 0.5 mL volumes at 45 °C, with 0.1 M KPi buffer at pH 5 or 9.5, containing 40 mM D-glucose, 10 mM pyruvate, 1 mM NAD^+^, 1 mg/mL lactate dehydrogenase (LDH), and 1 mg/mL glucose dehydrogenase (GDH). GDH-101 used for these experiments was supplied by Johnson Matthey. The reactions were incubated for 24 hours with shaking at 700 rpm. Triplicate reactions were performed for each condition and time point. Prior to initiating the reactions, the pH of the reaction mixtures was adjusted to the desired value (5 or 9.5) using the appropriate buffer. Control reactions were conducted with an empty vector (lyophilised powder) instead of GDH. At specific time points (0, 1.5, 3.5, 6, 20, and 24 hours), the reactions were stopped by adding 20 μL of concentrated TFA to precipitate the proteins. The samples were centrifuged at 4000 rpm for 20 minutes, and the supernatants were collected for analysis. The analysis was performed using an Agilent 1290 Infinity II UHPLC system equipped with an Aminex HPX-87H column (Bio-Rad) maintained at 35 °C. The samples were injected (20 μL) and eluted with 0.1% TFA (pH 2) at a flow rate of 0.6 mL/min for 25 minutes. The eluent was monitored by a UV detector at 210 nm. Standard solutions of pyruvate, lactate and gluconic acid (0-10 mM) were prepared and treated identically to the reaction samples for calibration purposes. The chromatograms were recorded, and data analysis was performed using OpenLab software (Agilent). The conversion of substrates was calculated by correlating the peak areas of the respective molecules to their concentrations using calibration curves (**Fig S19**).

### Experimental protocol for cascade 2

The second cascade reactions were conducted in 0.5 mL volumes at 45 °C, with 0.1 M KPi buffer at pH 5 or 9.5, containing 1 mM pyridoxal 5’-phosphate (PLP), 40 mM D-glucose, 50 mM D-alanine, 15 mM acetophenone, 1 mM NAD^+^, 1 mg/mL lactate dehydrogenase (LDH), 1 mg/mL glucose dehydrogenase (GDH), and 10 mg/mL transaminase (TA). GDH-101, RTA-25, RTA-57 and STA-14 used for these experiments were supplied by Johnson Matthey. The reactions were incubated for 24 hours with shaking at 700 rpm. Triplicate reactions were performed for each condition and each time point. Prior to initiating the reactions, the pH of the reaction mixtures was adjusted to the desired value (5 or 9.5) using the appropriate buffer. Control reactions were conducted with an empty vector (lyophilised powder) instead of the transaminase enzymes. At specific time points (0, 1.5, 3.5, 6, 20, and 24 hours), the reactions were quenched by adding 0.1 mL of 5 M NaOH and extracting twice with 0.6 mL of ethyl acetate. After each extraction, the samples were centrifuged at 4000 rpm for 5 minutes. For analysis, 0.7 mL of each sample was combined with 0.5 mL of ethyl acetate and 25 mM toluene (internal standard). Blank samples were prepared by mixing 1.2 mL of ethyl acetate and 25 mM toluene. The analysis was performed by injecting 1 μL of the sample onto an Agilent 7890B GC System equipped with an HP-5ms Ultra Inert GC column (15 m, 0.25 mm, 0.25 μm, 7-inch cage, 19091S-431UI). Helium was used as the carrier gas with a flow rate of 6.5 mL/min (**Table S8**). Standards for acetophenone (substrate, 0-8.5 mM) and (R/S)-α methylbenzylamine (MBA) (product, 0-15.6 mM) were prepared and treated as the reaction samples. The chromatograms were recorded, and data analysis was performed using OpenLab software. The conversion of substrates was calculated by correlating the peak areas of the respective molecules to their concentrations using the calibration curves (**Fig S19**).

