## Supporting Information for "Conditional language models enable the efficient design of proficient enzymes"

**Supporting Information Figures**

Figure S1-S19

**Supporting Information Tables**

Table S1-S10

### Supporting Information Figures

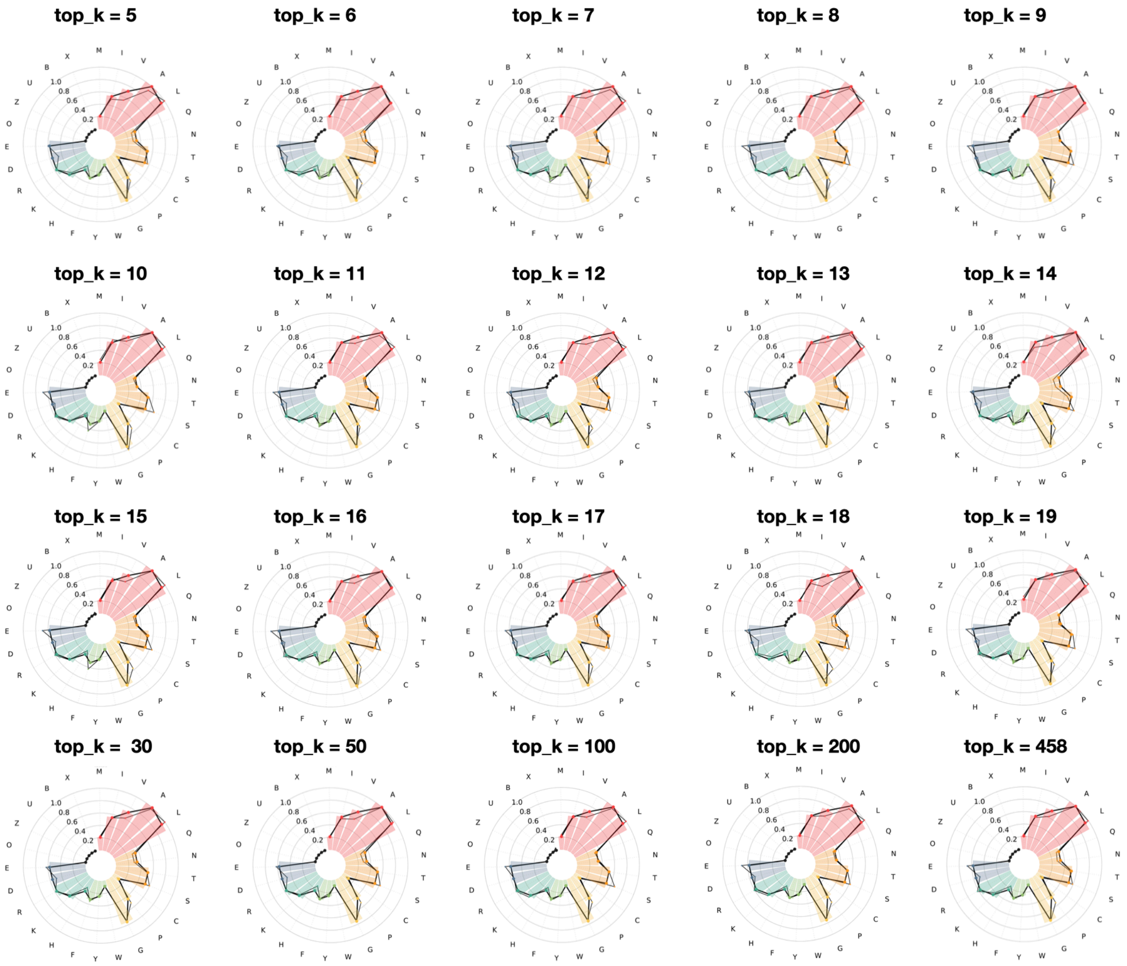


**Fig. S1:** Comparison of different parameter sets at recapitulating the natural sequences' amino acid propensity. The amino acids in the dataset are normalised (0,1) and shown in wider lines are the natural propensities. Different parameters approximate to different extents this distribution, with top_k = 9 being the closest.

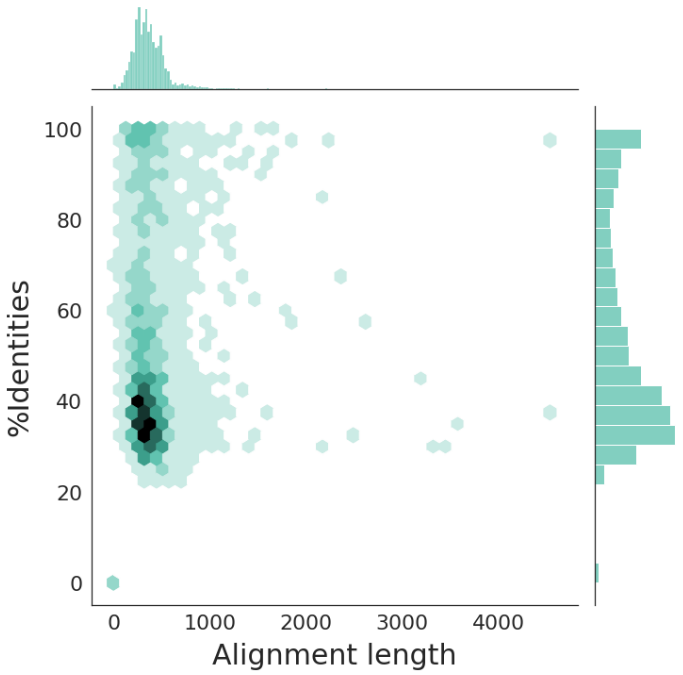


**Figure S2:** Identities and lengths for the best alignment found according to the E-value with BLASTp against the non-redundant protein sequence (nr) database.

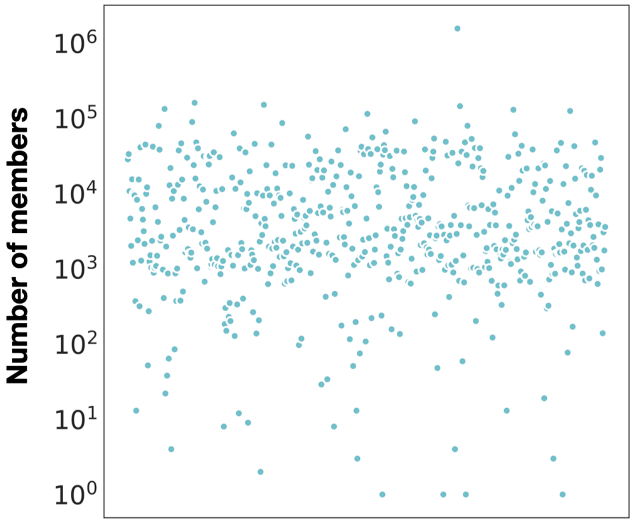


**Figure S3**: Number of members for each of the classes with hits over 90% in **Figure 2a** in the main text. The classes are shown in random order on the x-axis.

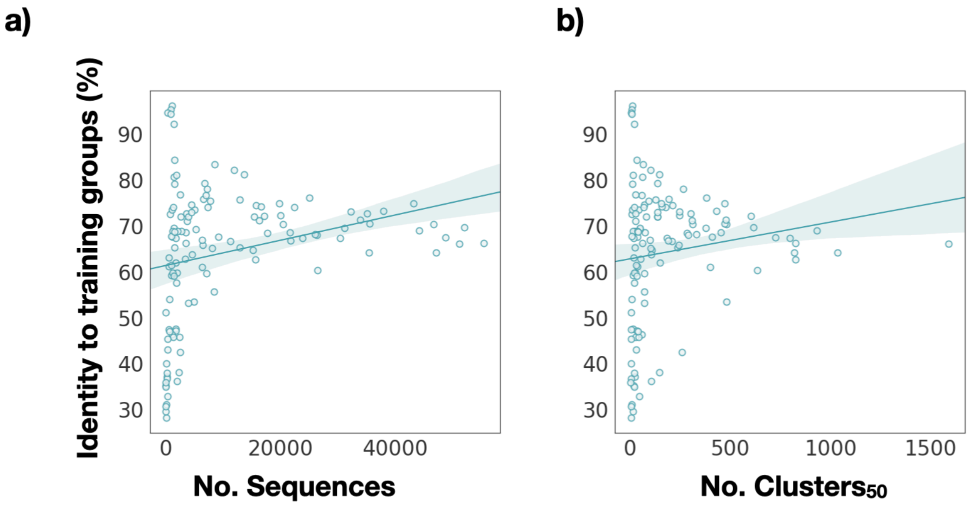


**Figure S4:** Mean identities for generated sequences to their training set EC classes as a function of **(a)** the number of sequences and **(b)** the number of clusters at 50% ($\text{Clusters}_{50}$). Each datapoint represents an EC class.

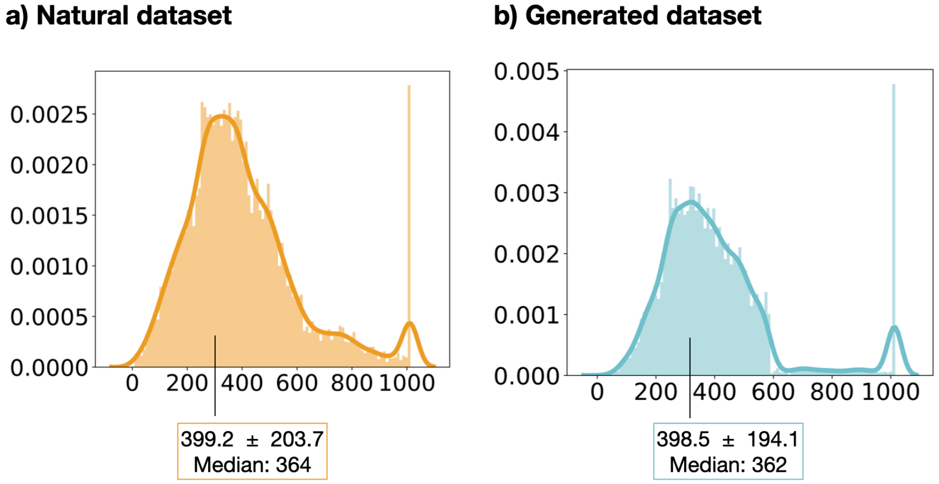


**Figure S5:** Length distribution for the natural **(a)** and generated **(b)** datasets.

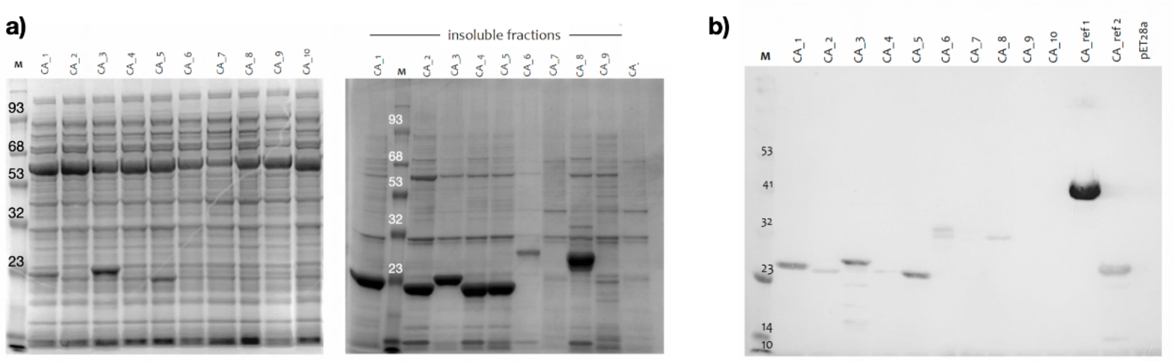


**Figure S6:** Protein overexpression for carbonic anhydrases CA1-10. **(a)** SDS-PAGE analysis of total protein (left) and the insoluble fraction (right) are shown. **(b)** Western blot analysis of soluble fractions for carbonic anhydrases CA1-CA10. Ca_ref2 corresponds in this case to the wild type sequence of *E. coli (*CA_ref, Table S2*)*. CA18 is a BaseGraph natural sequence included for testing purposes in this batch but not analysed in the manuscript. CA_ref1 corresponds to CA11, a generated fumarate hydratase (EC: 4.2.1.2) included as negative control.

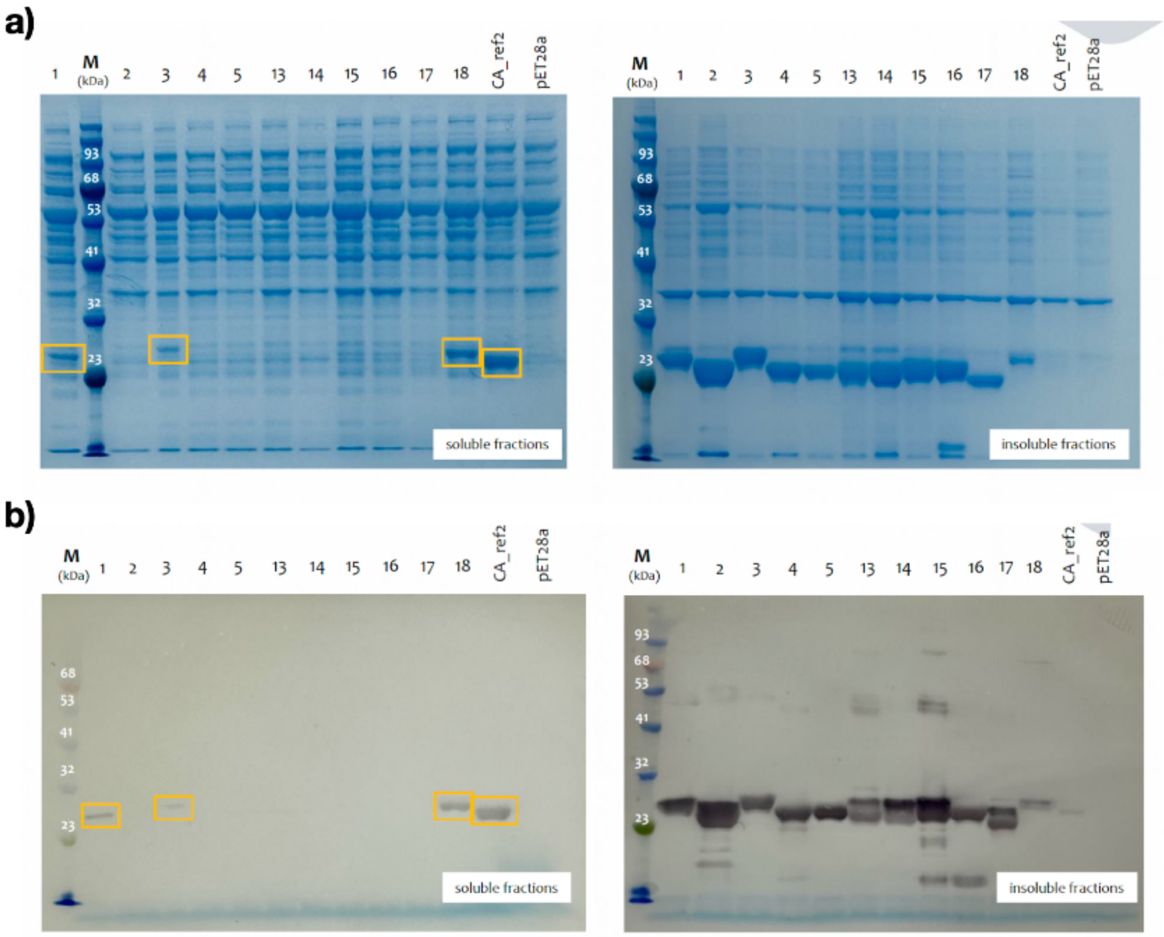


**Figure S7:** Protein over expression for carbonic anhydrases CA13 - CA17 along with previous analysed CA1 - CA5 (Fig S6). **(b)** Western blot analysis of soluble fractions for carbonic anhydrases CA13 - CA17 along with previous analysed CA1 - CA5 (Fig S6). Ca_ref2 corresponds in this case to the wild type sequence of *E. coli*. CA18 is a BaseGraph natural sequence included for testing purposes in this batch but not analysed in the manuscript.

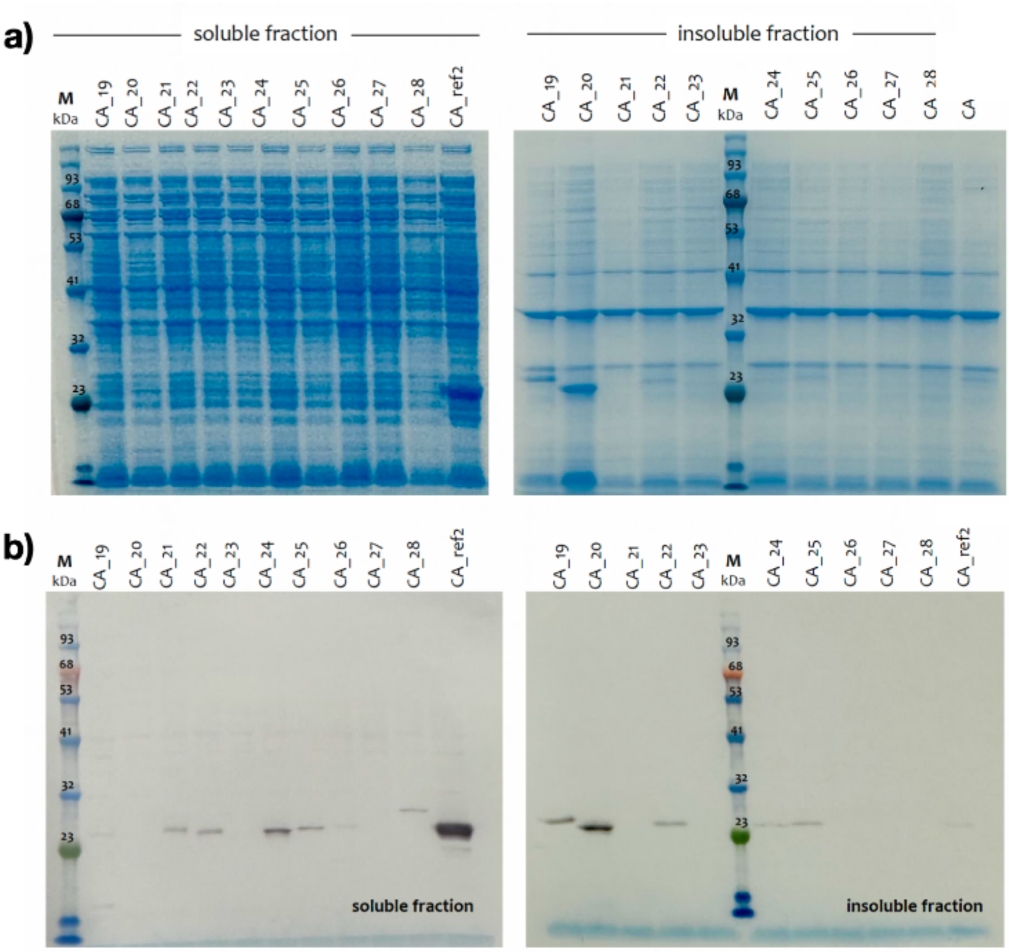


**Figure S8**: **(a)** Protein over expression for carbonic anhydrases CA19 - CA28. **(b)** Western blot analysis for carbonic anhydrases CA19 - CA28. Ca_ref2 corresponds in this case to the wild type sequence of *E. coli*.

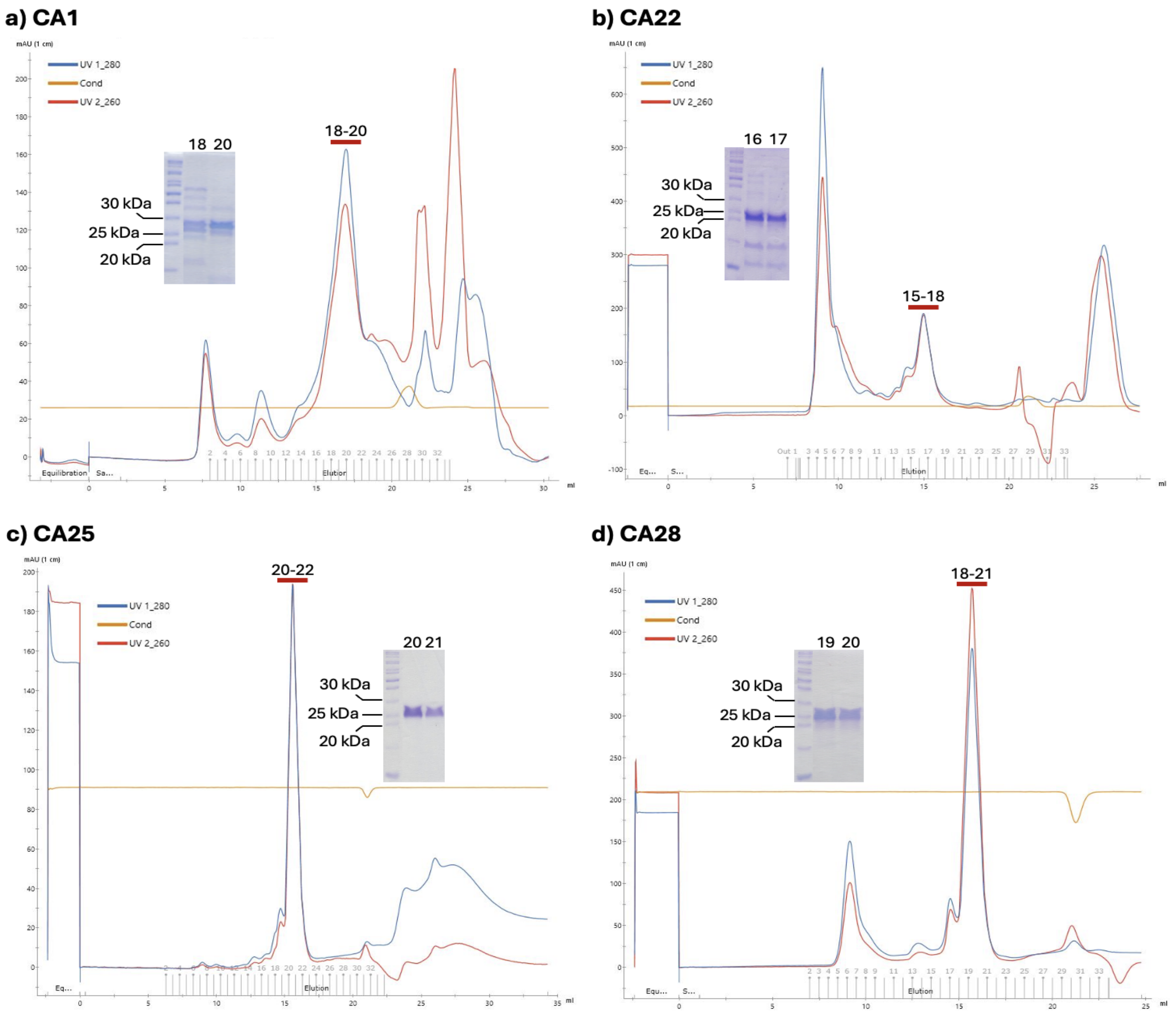


**Figure S9**: Size Exclusion Chromatography for **(a)** CA1, **(b)** CA22, **(c)** CA25, and **(d)** CA28. The SDS-PAGE at 15% acrylamide of the two main fractions of the peaks' maximum are shown. In all panels, UV280 and UV260 were normalised to 1 cm pathlength.

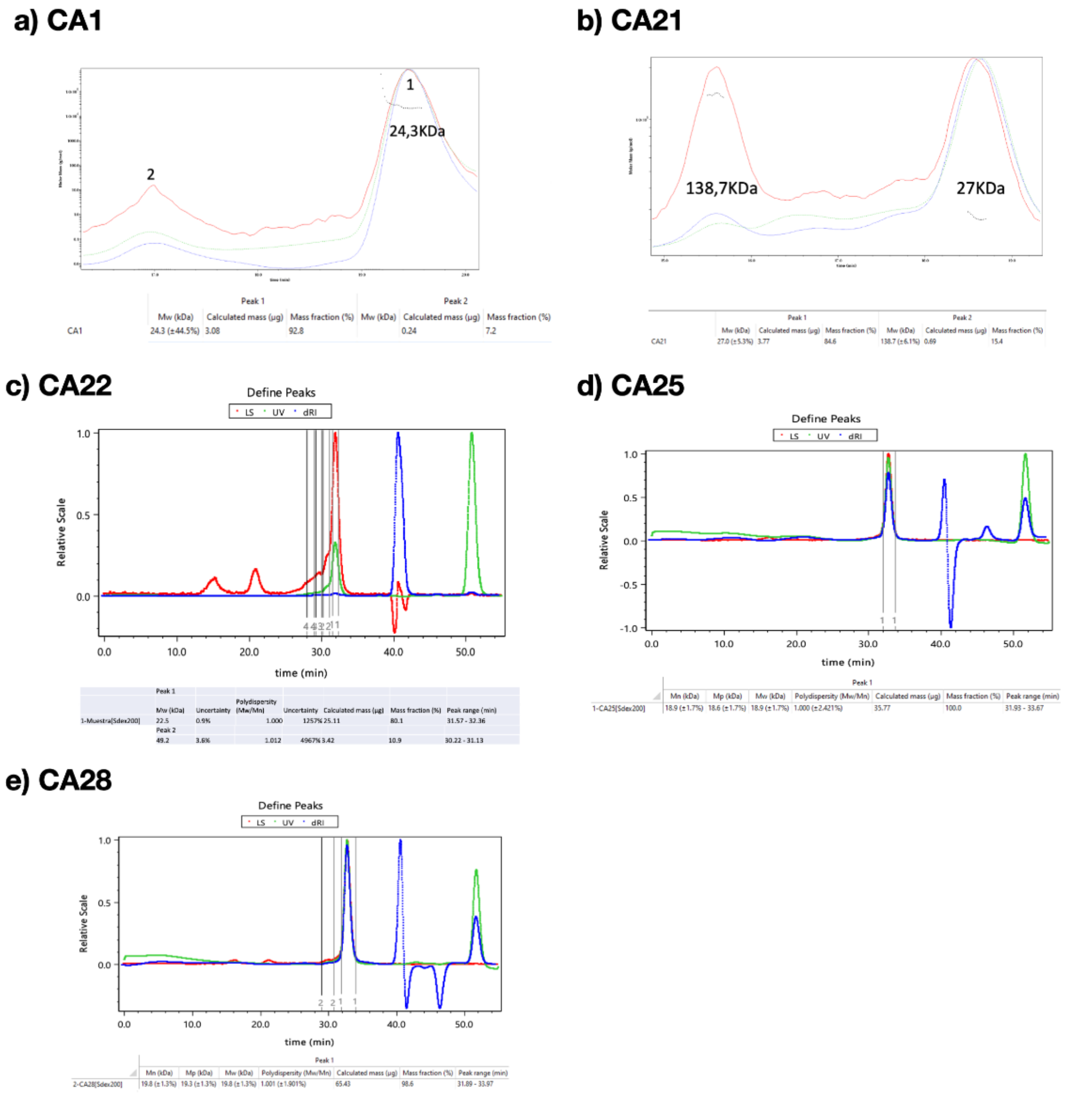


**a) CA1**

| | Peak 1 | | | Peak 2 | | |
|---|---|---|---|---|---|---|
| | Mw (kDa) | Calculated mass (µg) | Mass fraction (%) | Mw (kDa) | Calculated mass (µg) | Mass fraction (%) |
| CA1 | 24.3 (±44.5%) | 3.08 | 92.8 | | 0.24 | 7.2 |

**b) CA21**

| | Peak 1 | | | Peak 2 | | |
|---|---|---|---|---|---|---|
| | Mw (kDa) | Calculated mass (µg) | Mass fraction (%) | Mw (kDa) | Calculated mass (µg) | Mass fraction (%) |
| CA21 | 27.0 (±5.3%) | 3.77 | 84.6 | 138.7 (±6.1%) | 0.69 | 15.4 |

**c) CA22**

| | | | | | | | |
|---|---|---|---|---|---|---|---|
| | Peak 1 | | | | | | |
| | Mw (kDa) | Uncertainty | Polydispersity (Mw/Mn) | Uncertainty | Calculated mass (µg) | Mass fraction (%) | Peak range (min) |
| 1-Muestra[Sdex200] | 22.5 | 0.9% | 1.000 | 1257% | 25.11 | 80.1 | 31.57 - 32.36 |
| | Peak 2 | | | | | | |
| | 49.2 | 3.6% | 1.012 | 4967% | 3.42 | 10.9 | 30.22 - 31.13 |

**d) CA25**

| | Peak 1 | | | | | | |
|---|---|---|---|---|---|---|---|
| | Mn (kDa) | Mp (kDa) | Mw (kDa) | Polydispersity (Mw/Mn) | Calculated mass (µg) | Mass fraction (%) | Peak range (min) |
| 1-CA25[Sdex200] | 18.9 (±1.7%) | 18.6 (±1.7%) | 18.9 (±1.7%) | 1.000 (±2.421%) | 35.77 | 100.0 | 31.93 - 33.67 |

**e) CA28**

| | Peak 1 | | | | | | |
|---|---|---|---|---|---|---|---|
| | Mn (kDa) | Mp (kDa) | Mw (kDa) | Polydispersity (Mw/Mn) | Calculated mass (µg) | Mass fraction (%) | Peak range (min) |
| 2-CA28[Sdex200] | 19.8 (±1.3%) | 19.3 (±1.3%) | 19.8 (±1.3%) | 1.001 (±1.901%) | 65.43 | 98.6 | 31.89 - 33.97 |

**Figure S10:** SEC-MALs results for carbonic anhydrases (CA) **(a)** CA1, **(b)** CA21, **(c)** CA22, **(d)** CA25, and **(e)** CA28

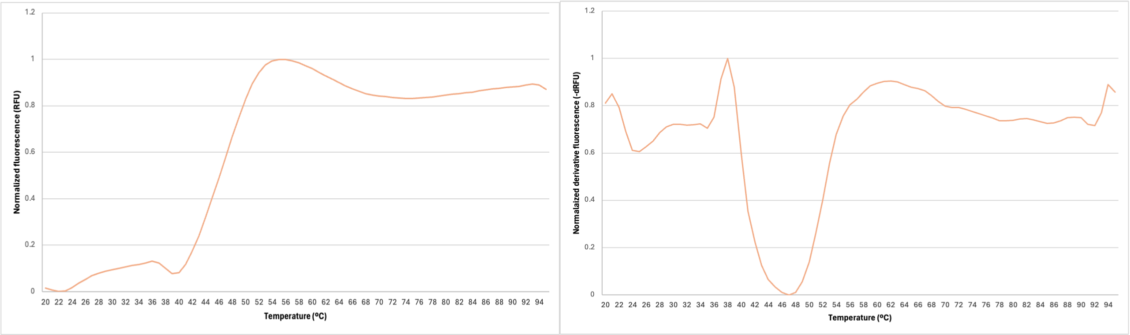


**Figure S11: DSF curves for carbonic anhydrase 22. Left,** The normalised relative fluorescence units (RFU) are plotted vs. the increased temperature (ºC), where the inflection point corresponds to the $T_m$ of CA22. **Right,** normalised derivative RFU vs. temperature (ºC), in which the inflection point from the left panel is the minimum of -dRFU, denoting its accurate value, 47 ºC.

## a) CA1

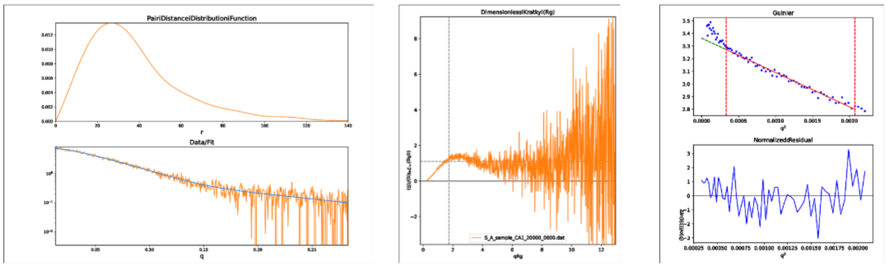


## b) CA22

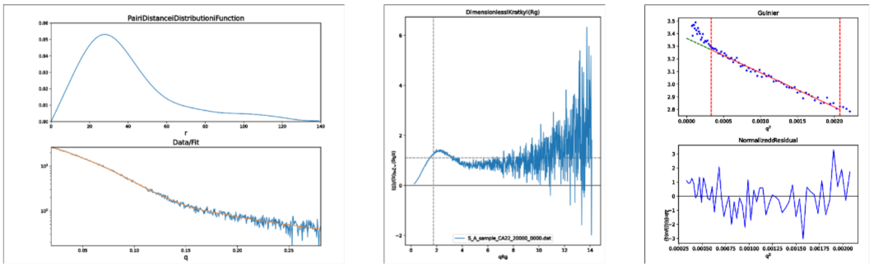


**Figure S12:** SAXS results for **(a)** CA1 and **(b)** CA22. From left to right, Pair-distance distribution function (P(r)), Kratky plot and Guinier analysis. For both enzymes, the $D_{max}$ calculated from the P(r) was 139 Å, and  the Kratky profile followed that tendency of a globular species. The Rg values calculated from the Guinier analysis for CA1 and CA22 were 28.23 ± 0.68 Å and 28.45 ± 0.22 Å, respectively.

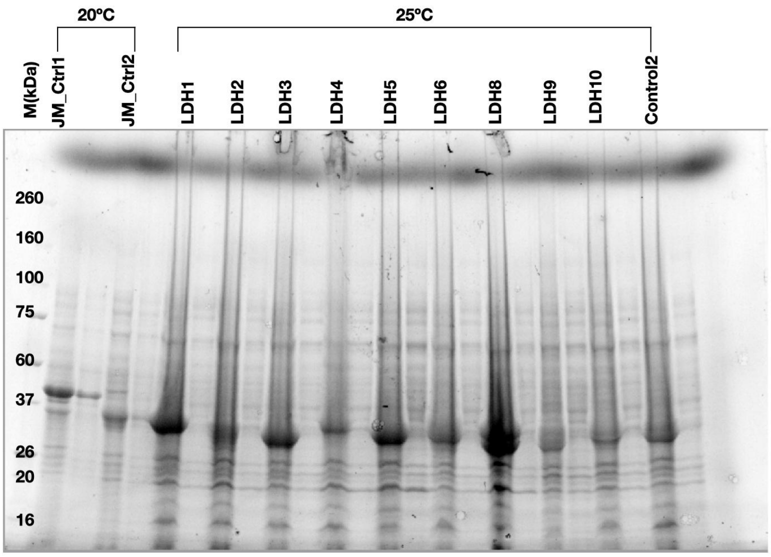


**Figure S13**: SDS-PAGE analysis of LDH1 - LDH6 and LDH8 - LDH10 at 25 °C, including Control2. Johnson Matthey proprietary LDH enzymes JM_Ctrl1 and JM_Ctrl2 at 20 °C were also included for testing but are not analysed in the manuscript. For each LDH the left lane corresponds to the total lysate and the right lane to the soluble fractions.

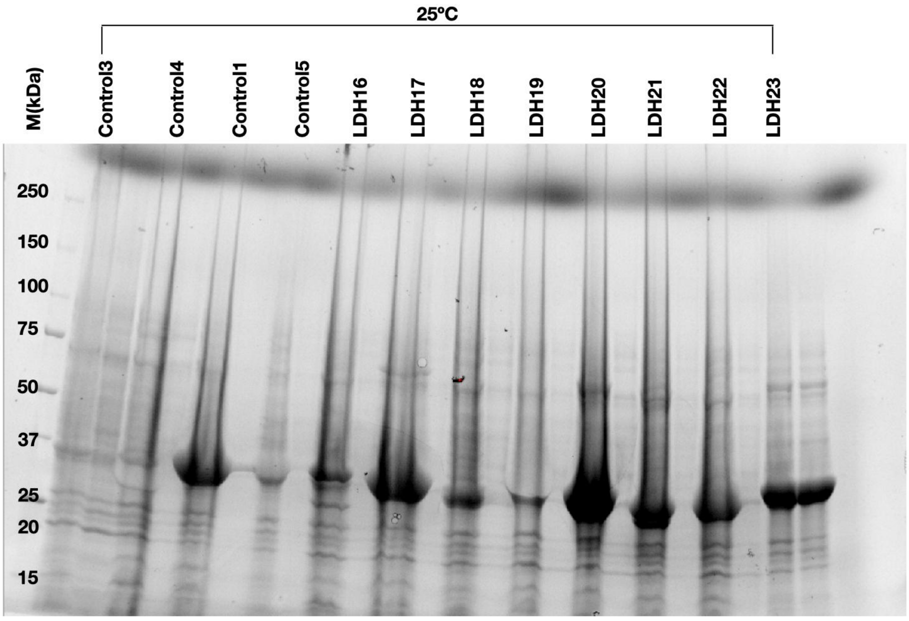


**Figure S14**: SDS-PAGE analysis of LDH16 - LDH23 at 25 °C, including Control1 and Control3. Control4 and Control5 were included for testing purposes but not analysed in the manuscript. For each LDH the left lane corresponds to the total lysate and the right lane to the soluble fractions.

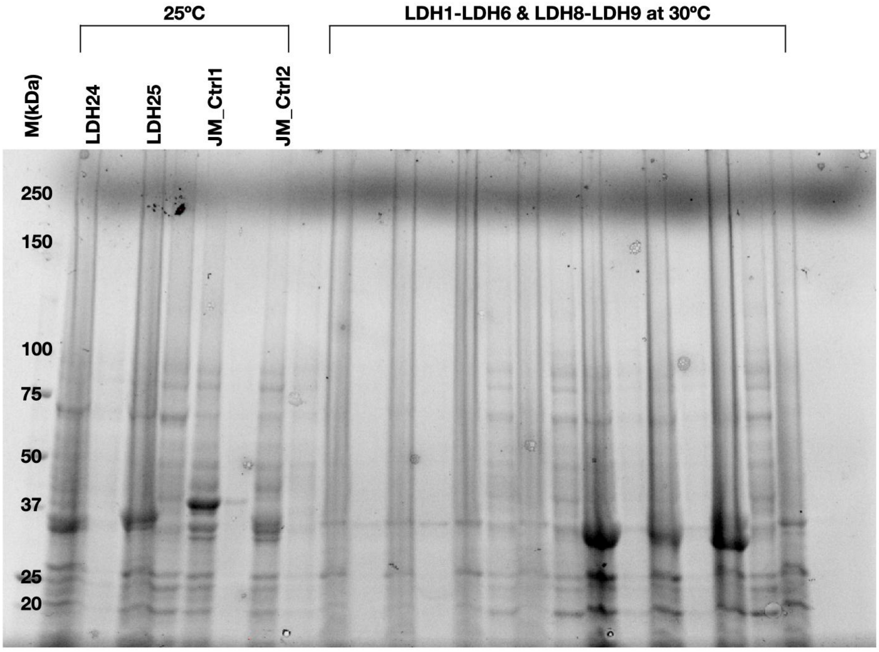


**Figure S15**: SDS-PAGE analysis of LDH24, LDH25 and JM_Ctrl1 and JM_Ctrl2 at 25 °C. For each LDH the left lane corresponds to the total lysate and the right lane to the soluble fractions. Additional lanes denote LDH1 - LDH6 and LDH8 - LDH9 at 30 °C.

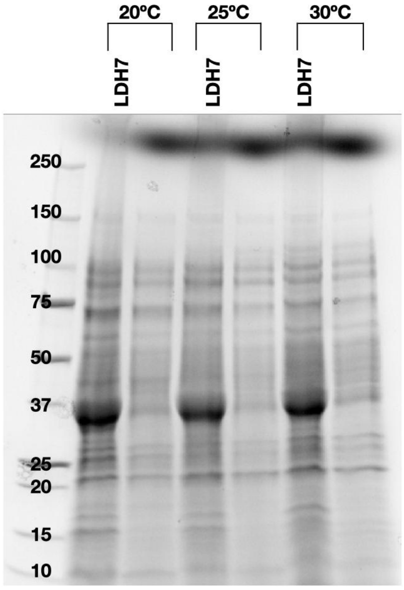


**Figure S16**: SDS-PAGE analysis of LDH7 at 20 °C, 25 °C and 30 °C. For each LDH the left lane corresponds to the total lysate and the right lane to the soluble fractions.

a) b)

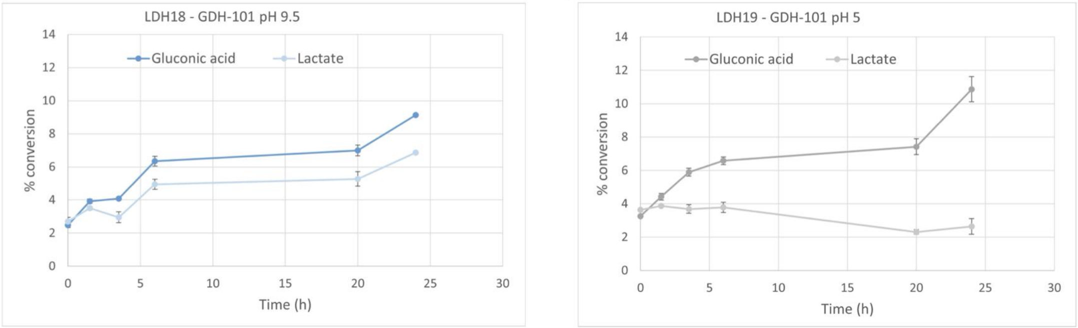


**Figure S17:** Rate of conversion of gluconic acid and lactate catalysed by **(a)** LDH18 and **(b)** LDH19 and GDH101 at pH 9.5 and pH 5, respectively for cascade 1.

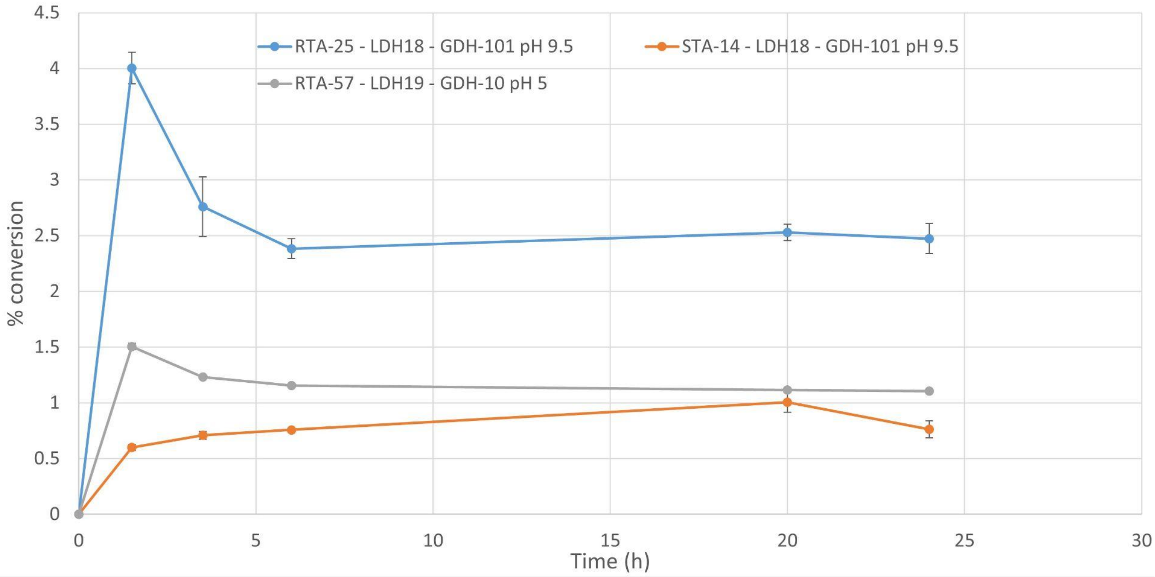


**Figure S18:** Rate of acetophenone to (R/S)-MBA conversion catalysed by R-transaminases RTA-25, RTA-57 and S-transaminase STA-14 in a multi-enzyme cascade reaction (cascade 2)**.** Reactions were performed at pH 5 and 9.5 as these were the optimum pH for LDH19 and LDH18, respectively (**Table S6**)

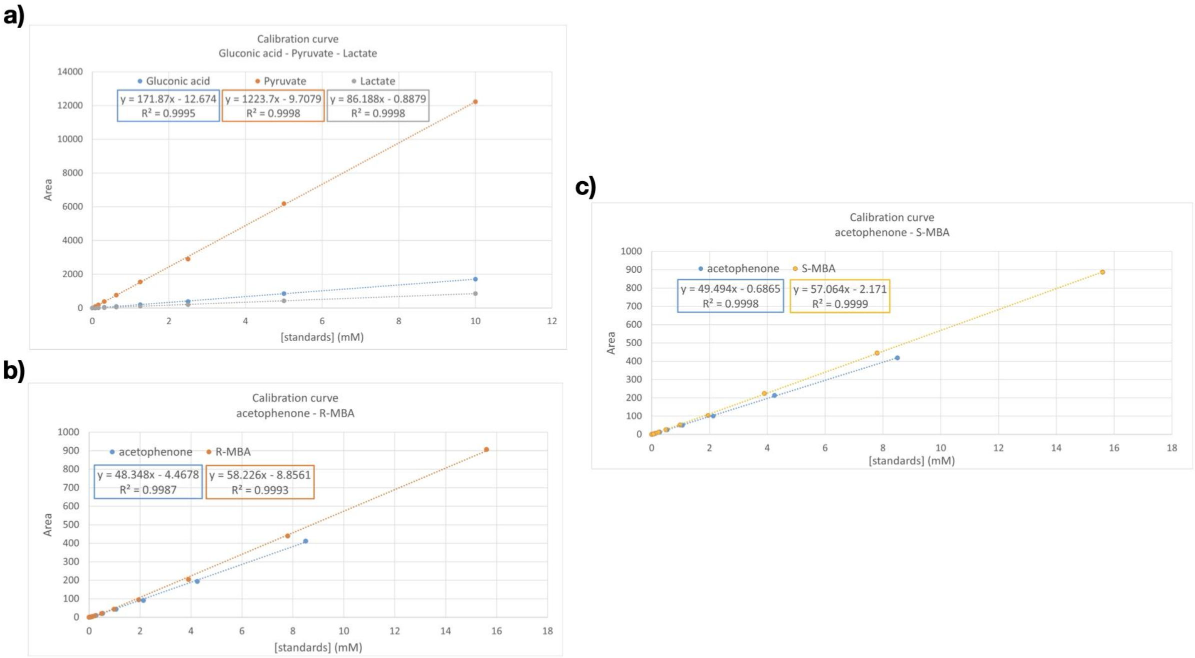


**Figure S19. (a)** Calibration curves of gluconic acid, pyruvate, and lactate, **(b)** acetophenone and R-MBA and **(c)** acetophenone and S-MBA

### Supporting Information Tables

**Table S1:** Labels used for generation in the invented and generated datasets.

| | Generated set | Invented (random) set |
|---|---|---|
| **EC label** | 2.1.1.334, 2.1.1.n8, 2.7.6.1, 2.1.3.10, 3.4.21.63, 2.7.1.189, 1.1.1.304, 2.1.1.166, 1.1.1.95, 3.4.21.109, 2.1.1.174, 2.4.1.315, 2.7.12.2, 3.6.5.5, 3.4.22.40, 3.6.3.1, 3.1.11.5, 2.1.1.131, 2.1.2.1, 2.7.11.18 | 9.9.9.9, 99.9.4.3, 9.9.90.0, 99.88.77.7, 9.9.3.3, 9.7.7.99, 9.6.6.66, 9.6.6.3, 9.6.5.6, 9.4.44.44, 9.11.11.1, 8.9.99, 8.8.8.88, 88.8.2.2, 8.5.5.5, 8.4.455.5, 8.2.2.2, 7.7.7.7, 33.44.22.1, 0.0.0.9 |

**Table S2. Summary of properties for the tested carbonic anhydrases.** Enzyme ID: Original naming scheme; Testing ID: Naming followed through the manuscript. CA subtype: carbonic anhydrase subtype (alpha or beta); Sequence identity: best hit with BLAST against the nr database; Length: number of amino acids; pLDDT: ESMFold average over all residues; TM score: TM align result against PDB 1CA2 (alpha) or 1DDZ (beta); HSASA: hydrophobic surface accessible solvent area; Ideal surface * 1.7 computing ideal surface of a protein following Dill et al.[1]; Net charge: total charge computed with localcider[2]; IP: isoelectric point computed with biopython; ProteinMPNN: ProteinMPNN score obtained for the ESMFold prediction and the corresponding sequence. The second batch is depicted in grey.

| Testing ID | CA subtype | Sequence identity | Length | pLDDT | TM score | HSASA | Ideal surface * 1.7 | Net charge | IP | Protein MPNN |
|---|---|---|---|---|---|---|---|---|---|---|
| CA1 | beta | 46.4 | 209 | 91 | 0.84 | 7725.09 | 7065.6 | 2 | 8.31 | 1.84 |
| CA2 | beta | 42.6 | 216 | 91 | 0.80 | 7749.27 | 7250.5 | -2 | 6.45 | 1.85 |
| CA3 | beta | 43.2 | 212 | 92 | 0.81 | 7080.12 | 7145 | -11 | 5.36 | 1.91 |
| CA4 | beta | 42.4 | 202 | 92 | 0.84 | 6934.58 | 6879.4 | -12 | 4.92 | 1.84 |
| CA5 | beta | 48.8 | 202 | 91 | 0.84 | 6860.24 | 6932.7 | -13 | 4.96 | 1.79 |
| CA6 | alpha | 43.3 | 247 | 84 | 0.83 | 7172.73 | 8054.3 | -11 | 5.27 | 1.90 |
| CA7 | alpha | 39 | 266 | 83 | 0.86 | 6972.84 | 8536.1 | -18 | 4.64 | 1.88 |
| CA8 | alpha | 45.7 | 246 | 85 | 0.80 | 7831.41 | 8028.8 | -4 | 5.02 | 1.79 |
| CA9 | alpha | 51.3 | 255 | 89 | 0.83 | 8546.19 | 8258.1 | 10 | 9.52 | 1.76 |
| CA10 | alpha | 43.8 | 249 | 86 | 0.82 | 7396.23 | 8105.4 | -9 | 5.45 | 1.80 |
| CA11 (fumarate_hydratase) | - | 99.2 | 399 | - | - | - | - | - | 6.02 | - |
| CA_ref (Escherichia coli (strain K12) | beta | - | 219 | 94 | 0.86 | 7537.02 | 7329.3 | 0 | 6.99 | 1.59 |
| CA13 | beta | 46.1 | 199 | 92 | 0.81 | 6945.78 | 6799.2 | 3 | 8.59 | 1.73 |
| CA14 | beta | 48.7 | 208 | 91 | 0.83 | 7204.80 | 7039.1 | -4 | 6.24 | 1.88 |
| CA15 | beta | 48 | 210 | 92 | 0.80 | 6675.58 | 7092.1 | -8 | 5.54 | 1.88 |

| Testing ID | CA subtype | Sequence identity | Length | pLDDT | TM score | HSASA | Ideal surface * 1.7 | Net charge | IP | Protein MPNN |
|---|---|---|---|---|---|---|---|---|---|---|
| CA16 | beta | 48.5 | 211 | 93 | 0.81 | 6933.89 | 7118.6 | -11 | 5.05 | 1.74 |
| CA17 | beta | 47.5 | 209 | 90 | 0.81 | 7015.79 | 7065.6 | -6 | 5.32 | 1.77 |
| CA19 | beta | 50.8 | 211 | 90 | 0.78 | 7093.52 | 7118.6 | -3 | 5.83 | 1.80 |
| CA20 | beta | 43.6 | 205 | 90 | 0.79 | 6911.32 | 6959.4 | -12 | 4.68 | 1.83 |
| CA21 | beta | 50 | 215 | 82 | 0.78 | 7154.84 | 7224.1 | -10 | 5.09 | 2.07 |
| CA22 | beta | 47.6 | 208 | 93 | 0.80 | 6895.87 | 7039.1 | -7 | 5.47 | 1.81 |
| CA23 | beta | 44.9 | 201 | 92 | 0.80 | 6670.70 | 6852.7 | -4 | 5.71 | 1.69 |
| CA24 | beta | 42.9 | 207 | 89 | 0.75 | 7270.58 | 7012.5 | -14 | 8.79 | 1.71 |
| CA25 | beta | 50.9 | 206 | 88 | 0.74 | 6809.96 | 6986 | -6 | 5.30 | 1.91 |
| CA26 | beta | 48.8 | 210 | 84 | 0.26 | 7027.54 | 7092.1 | -9 | 5.21 | 1.77 |
| CA27 | beta | 43.3 | 211 | 89 | 0.79 | 5737.84 | 7118.6 | -12 | 4.94 | 1.68 |
| CA28 | beta | 40.5 | 225 | 87 | 0.72 | 8432.96 | 7486.3 | -12 | 4.94 | 1.91 |

**Table S3: Summary of solubility of carbonic anhydrases**

| Carbonic anhydrase # | Result |
|---|---|
| **CA1** | Expressed, Faint band soluble fraction (most protein found in the insoluble fraction) |
| **CA2** | Expressed, insoluble |
| **CA3** | Expressed, Faint band soluble fraction (most protein found in the insoluble fraction) |
| **CA4** | Expressed, insoluble |
| **CA5** | Expressed, Faint band soluble fraction (most protein found in the insoluble fraction) |
| **CA6** | Expressed, insoluble |
| **CA7** | Not expressed |
| **CA8** | Expressed, insoluble |
| **CA9** | Not expressed |
| **CA10** | Not expressed |
| **CA13** | Expressed, insoluble |
| **CA14** | Expressed, insoluble |
| **CA15** | Expressed, insoluble |
| **CA16** | Expressed, insoluble |
| **CA17** | Expressed, insoluble |
| **CA19** | Expressed, insoluble |
| **CA20** | Expressed, insoluble |
| **CA21** | Expressed, soluble |
| **CA22** | Expressed, soluble |
| **CA24** | Expressed, soluble |
| **CA25** | Expressed, soluble |
| **CA26** | Expressed, soluble |
| **CA27** | Not expressed |
| **CA28** | Expressed, soluble |

**Table S4**: **Summary of properties for the tested Lactate dehydrogenases.** Testing ID: Naming followed through the manuscript. Sequence identity: best hit with BLAST against the nr database; ppl: perplexity; Length: number of amino acids; pLDDT: ESMFold average over all residues; TM score: TM align result against (PDB: 1LDB); HSASA: hydrophobic surface accessible solvent area; Ideal surface * 1.7 computing ideal surface of a protein following Dill et al.[1]; Net charge: total charge computed with localcider[2]; IP: isoelectric point computed with biopython; ProteinMPNN: ProteinMPNN score obtained for the ESMFold prediction and the corresponding sequence.

| Testing ID | Length | Ppl | pLDDT | TM | HSASA ($Å^2$) | Ideal surface * 1.7 | Net charge | Isoelectric point | ProteinMPNN |
|---|---|---|---|---|---|---|---|---|---|
| Control2 | 310 | | 91 | 0.96 | 7454.07 | 10011.9 | -5 | 5.83 | 1.38 |
| Control3 | 314 | | 92 | 0.95 | 8285.47 | 9867.2 | -11 | 5.47 | 1.53 |
| LDH1 | 320 | 1.29 | 93 | 0.94 | 8041.76 | 9794.6 | -9 | 5.38 | 1.46 |
| LDH2 | 317 | 1.37 | 92 | 0.96 | 8015.57 | 9794.6 | -10 | 5.56 | 1.44 |
| LDH3 | 318 | 1.34 | 93 | 0.94 | 7832.91 | 9818.8 | -12 | 5.11 | 1.49 |
| LDH4 | 319 | 1.46 | 90 | 0.94 | 8440.48 | 9843.0 | -8 | 5.70 | 1.50 |
| LDH5 | 318 | 1.50 | 91 | 0.94 | 8682.30 | 9818.8 | -8 | 5.39 | 1.48 |
| LDH6 | 326 | 1.41 | 92 | 0.92 | 8895.64 | 10011.9 | -9 | 5.26 | 1.48 |
| LDH7 | 318 | 1.44 | 92 | 0.95 | 8463.94 | 9818.8 | -10 | 5.46 | 1.47 |
| LDH8 | 318 | 1.50 | 90 | 0.96 | 8045.01 | 9818.8 | -9 | 5.47 | 1.47 |
| LDH9 | 329 | 1.47 | 90 | 0.91 | 9197.32 | 10084.1 | -9 | 5.42 | 1.46 |
| LDH10 | 320 | 1.30 | 90 | 0.95 | 8559.80 | 9867.2 | -9 | 5.17 | 1.51 |
| LDH16 | 318 | 1.72 | 91 | 0.95 | 8236.48 | 9818.8 | -10 | 5.36 | 1.48 |
| LDH17 | 321 | 1.14 | 91 | 0.95 | 8434.39 | 9891.3 | -9 | 5.34 | 1.48 |
| LDH18 | 334 | 1.18 | 86 | 0.96 | 9579.23 | 10204.0 | -3 | 6.25 | 1.65 |
| LDH19 | 331 | 4.54 | 31 | 0.42 | 13593.83 | 10132.1 | -19 | 4.66 | 2.31 |
| LDH20 | 323 | 1.10 | 91 | 0.96 | 8563.27 | 9939.6 | -12 | 5.05 | 1.50 |
| LDH21 | 317 | 2.36 | 92 | 0.96 | 8454.61 | 9794.6 | -8 | 5.45 | 1.50 |
| LDH22 | 316 | 1.82 | 86 | 0.96 | 7763.70 | 9770.3 | -127 | 4.88 | 1.55 |
| LDH23 | 328 | 1.70 | 90 | 0.92 | 8677.80 | 10060.1 | -8 | 5.33 | 1.48 |
| LDH24 | 335 | 1.94 | 77 | 0.94 | 10240.72 | 10228.0 | 3 | 8.59 | 1.86 |
| LDH25 | 309 | 2.50 | 90 | 0.93 | 7843.92 | 9600.2 | 1 | 7.78 | 1.69 |

**Table S5: Summary of solubility of Lactate dehydrogenases**

| Lactate dehydrogenase # | Result |
|---|---|
| **LDH1** | Expressed, mostly insoluble |
| **LDH2** | Expressed, mostly insoluble |
| **LDH3** | Expressed, mostly insoluble |
| **LDH4** | Expressed, mostly insoluble |
| **LDH5** | Expressed, mostly insoluble |
| **LDH6** | Expressed, mostly insoluble |
| **LDH7** | Expressed, mostly insoluble |
| **LDH8** | Expressed, soluble |
| **LDH9** | Expressed, mostly insoluble |
| **LDH10** | Expressed, mostly insoluble |
| **LDH16** | Expressed, mostly insoluble |
| **LDH17** | Expressed, mostly insoluble |
| **LDH18** | Expressed, soluble |
| **LDH19** | Expressed, soluble |
| **LDH20** | Expressed, soluble |
| **LDH21** | Expressed, mostly insoluble |
| **LDH22** | Expressed, mostly insoluble |
| **LDH23** | Expressed, highly soluble |
| **LDH24** | Expressed, soluble |
| **LDH25** | Expressed, mostly insoluble |

**Table S6: Sequences of the tested carbonic anhydrases and lactate dehydrogenases**

| CA ID | Sequence |
|---|---|
| CA1 | MLKRLIQGFESFKRDYYTKHRKLFEELKKGQKPTVALFSCSDSRINPNQITQSNLGEIFIIR<br>NAGNLVPPYDSSNYGTSAAIEFAVCSLGVENIIVLGHSHCGGIEAILKEEGEKSFNKLVNK<br>WMKIADTAKDLLKNHLPDEDQLRKAAEINVRTQLDHLQSYPFISKKLKENELSIHAWVYHI<br>QTGSVFTFSQEKSGYFELEQKKINE |
| CA2 | MRKFEQLVEGYQTFRSTYFAHREALFNELSKGQRPATLIISCSDSRVDPNIILHTQPGDMFI<br>ARNIGNMIPPFSTASQYHGASTAIEFSIRSLKIKEIIILGHSGCGALKGILTPEPSANSYHYVE<br>NWISLAAPAREKLEALPSLQDKEKQIRLCEQHSIVIGVNHLMTFPFVKQRLADNQLNLHGL<br>WTDIGAGTLEWLESTNSTFKTVLNELPAD |
| CA3 | MNKKDIEKLISGFKRFQADYYPSRKEMVRELAAGQHPETLMISCVDSRIAPEHIAAGSPGD<br>IYVIRNAGNIIPEFETASGGESATVEFAVGALDIPHILVMGHSGCGAIKRFIEDTQEESSNSD<br>HIHDWLSLCAPVRELVAEHYGEAREAARVRAVKMRNVMAQLANLRSFPFIENRASEEKL<br>QLHGWHYDLATGEVWLFDHESDQFIRTEA |
| CA4 | MKKLIQGFVQFRTHHFSEQPAIYQRICDGQTPDVLVIACSDSRVDPAILLGCEPGDLLVTR<br>NVGNLVPPSDSDPSAAGIQYSIEYLQVKHIIVVGHASCGGVKALLEPQQAPSAENDFIGQ<br>WIRLAEPCHRTVLESPSQLSADTLHDLTVQEAVENLSTHTVLESRLQSGEMAVHGWLFSL<br>STGEVRALDQSANTFSEIPAA |
| CA5 | MKNLIQGIHRFQKDVFTEESALYRQLVSGQKPEVLYITCSDSRIDPHIMTQGDLFLVRNAG<br>NIIPPHGNTGEGIGSAVEFAVSGLNIRQVIVCGHSNCGAMQAILSPSSVKTMPALEQWLEH<br>IEDTVRVVRQHCHESLDEAARIKELVSQENVLQEVEHLMAYPVVEGIIKESKLQLQAWFYD<br>ISTGEVYVYDAETGTFQPIED |
| CA6 | MLRSFPIVAAGLISITSVHSADWGYKGNEDAPEHWGETSPEFSTCGNDQQSPINIKNATS<br>DTNLADSVKVGFEPTNYSNTSITNNSHTVIVGTNTIENGTSVNGHTYEFKQVHFHAPSEK<br>LEGKNFALEAHFVNSAKDGSLGVLAVFFQKGSANSSLTKLMTQINDNKEAIELGSLDPKE |

| | |
|---|---|
| | LLPDSTEYYHYKGSLTTPSCEEGIKWHVYKNTETISAKEASEISKIMGNNNRPVQEMNGR<br>TINLSTE |
| CA7 | MRSLVQLSALLVIGASHWTYCDLSPSTWGGLCDKTTEKSNPINIVTADTNTNENVDFTLNS<br>FDNITIINLQNNGHSVQVNLAEEVVLGGGLEIEGENYQLVQLHFHWGDEENPGSEHSLDG<br>KTYPMEMHLVNFKEGFFTDDIAKKPENKSYSVIGFLFEVGNENNAFQPISQKVSNITDKTE<br>STNIAIDIETLLPSNSQFYRYTGSLTTPSCAEVVSWYVLPDPVTLSEKQIKKLEQLTESDKQ<br>LEGNYRSTQPIHQRTVTANSAN |
| CA8 | MKRLFSGALVTVCAAPLAASAGPHWSYEGEAGPENWGSLSEDYTACEAGSQQSPIDVT<br>EAIDDRVAPIKLNYSGASTLINNGHTIQAAFAQGSTLIVGGTLKTIAGGGLAGTYEARQFHF<br>HAPSEHMVDGKRFPIEAHFIHADRQDAGLAVIGVMLEVGAENPAIAAAWDALPAQVGELA<br>AVPGTDRFYKYNGSLTTPACDEVVTWQILRDPIEVSREQIEKFTTVMGEGNARPVQPLNA<br>RIVLSR |
| CA9 | QSILKHAVPAIALVLCTFALAPSAAHGKEWGYEGASGPKHWGQLDKDNFACSQGKNQSP<br>IDIVSDKSVKSTLKPIRRNYKPSNATITNNGHTIAVEYAAGKELLINGKTYELKQFHFHTPSE<br>HLVNGKSYPLEVHFVHVAETGELAVVGVMFEQGSTNPALQSMWAKLPAKTTGAGQLQA<br>AFNAEGLLPTNRDSFYRYKGSLTTPPCAEGVRWTVLKQPATVAAERMAKFNKTMAHPNA<br>RPLQPVNARPVKSGA |
| CA10 | MKSFILALVFIPVCAYAADWGYQTEHAPENWSAISAENSRCGEGRQSPVNIDTSVKVVPL<br>KSALNLTYGPTILHSITNTGDTISAQWKGSAGKLVIDGNKYQLAQFHFHTPSENHINGTSY<br>PMELHLVHKNAESDLLVLGVLVEPTEGSEHSLSQLFAEVPNAEGALVKEKELLTLPLNVKQ<br>LLPKNRSYYHFNGSLTTPPCTQGVQWYVLAAEPMEISQDQIARFRSLMNNEENNPLQPL<br>NSRVVMED |
| CA13 | MNKIENLIKGFKNFQREYFSEDTQLFKTLAKGQKPHTLIVTCADSRIETALLANAGPGDIFV<br>ARNIGNLVPPHTIEGGGSAAALEFGIQALDCPEIIVCGHSHCGAMKGLANPANLKALPNVR<br>DWLALAAPCRRLIKTKYPELDDAERVDAVAQRNVIAQTGLLNNLLTYPWVKKRVEQGELKI<br>HALFFDIASGTLHKL |
| CA14 | MKKLFEGLLHFQKHFYDNDDQLFSQLKNGQTPRVLLIACTDSRIDPNLITHTMPGDLFVMR<br>NMGNLIPTHGLGGRFSSISAAIEYGLLNLQVEDVIVCGHSDCGAMNALHTHPKLKNLLPVL<br>AAWLKLAEPTQRITAQHPGSTDEEIRHVNIANVLEQMEHLRTYPLVGELLQDGQLRLYGW<br>YYIIETGHIYNYNDNRKAFKLIENEK |
| CA15 | MTSPLPPRLAQGYHSFVQERLADHERLFEKLAAGQKPRTLLIACCDSRVDPAMVLGCDP<br>GDIFVLRNLGNFVPPYEGADHQHGTSAAIEFAVQVLEVAQIIILGHSQCGGITHLSGEIETLP<br>NVATWLGHGRQALERIRDGHASDDLVTPEERQSCEKRSIRQSLDNLVTFPRVQERLQNG<br>QLKIHGWYFDLDDGGVMGYDPQAGRFTTI |
| CA16 | MRDIERLVAGFKRFQDHYYTSHPDLFGQLSQGQNPQAFVVTCADSRVEPALLTNCEPGE<br>LFNLRNITNVLPPFERDDKEHALHASIAFAVEELHVKKIVVLAHSNCGGIRALMQGSDSGE<br>GDLSYVNEWVAIAEPAKQVVRNNYPDLEPDQLRRLCERESIMVSLENLMTFPFIAERVAQ<br>GELTIHAWYYNIAKGELLNLNQQTSEMTVIA |
| CA17 | MSTDIKEALERLHDGYRAFLEGQFPRYQAVFTELSQQRPEILVLACCDSRVDPAVITECGP<br>GDIFLIRNVASLVPPCATAAGGASGTRAAIEFAVLALQVNKIIVLGHSNCGGIRALYDGAVT<br>GSDFLQSWVSLVRQASESVAGDPDPEGLVAQAQHAAVRAAIARLRDYPVLAARREAGAV<br>RLHGWHFDLATGEIWALDGAGGFFPIG |
| CA19 | MNDLARFVRGFRHFQQSYYRADERGLYTSLAQSQRPAMLIACADSRADPGVVFEAGPG<br>EMFVARIAGNIVPPYQPDGQFLGVSAALEFAVNILKVGSIVVMGHSLCGGCKALMALDPQ<br>PAQEPSEDFISQWMQVAQAARDRIIEARGPAERRQVEQLEREAVRRSLRNLMTFPYVRA<br>RVESGEIALHGWHFDIETGGVDWLSRSADGFLPV |
| CA20 | MPLSDLFARNREWVEQTLKKDTDYFSKLSTGQKPRITIIACCDSRVHVTALGLGPGVAFTT<br>RNVAALIPPYQTGQGSALEVTTNAALEFALTALEVNHAVVVGHGCCGGIRAFVDEASKGK<br>EDEADEEPANLADAWLKLLDEVSARVEAEYAEGRDPAFVTELEVQRTVANLLGFPWIEQA<br>ISDRGLLRIDGVIFDVAGGRLSEL |
| CA21 | MASETYEDAIKGLKNLLSEKGEPLPVAAVNVVRKLESDPQYFDEMKQRFDTGQRPQALFI<br>SCADSRVCPSHIVGFHSGEVFMVRNVANMVPSDDTSASSAIEFAVTQLKVKNILVMGHCK<br>CDGLSAMNTDELHSNMLEPWVKEASSTTLKLVNGEDLSFEDQCQKCEKESVKRSIQQLS |

| | |
|---|---|
| | HLRSYPFVKEAQANDQLAIHGGYYDFIEGSFETWER |
| CA22 | MQELVEGFKSFQRDAYPQLSRFQKELISRGQSPQTMIVACCDSRADPTMILNCDPGDIFVIRNVANLISPYHPHADGVHAASAALEFAVQALKVKHIIILGHAGCGGIAAFAQSQDLPDSDDYISAWMKILEPARQVTLKALGNDRTAADRACELESVRVSLQNILTFSWIRERVERGELRIHGWWFDVSAGEVLAYSAASHQFEPID |
| CA23 | MSADLKQRIQSGYTRFRETGLDSQASLYSNLATAHQPTAVMISCVDSRALPEVIFDAGLGDMFVVRLAGNIVAPSQSGGVTASVEYAVSQVHGTPLVVVLGHSHCGAISAIKNDILNGAPLPGYVGRLMDHIEPAMRQLPDESVDNAARANALLTANAVARSIETIRAQSPLIKAAVDSKKLAVVGALYRLDTGEITFYEE |
| CA24 | MSTDLTDAPEPVSLAAFLTANDEFVRQAAQHPAGDTGFGLSRTGVFPDRVIVSCSDAAPAETVVNARPGEAFEVRVAGNRILDESVVASIEYAVTALGTELLLVLGHEGCGAVKTAAEQLAEGAVESGSFLAHLRPVVEAVRASGDLRETTEEERRRHADAVIRTSIALSVEHLRTFPWVQAAVTDGTVTVLTGHFEISDATLARLG |
| CA25 | MTTNELQKRIAKGFDKYQHQYFDESPDLYQKLSEKQQPKAMVVTCCDSRADPATQIMGMGPGNAFIVRNVANLVPVVEPGTGQHGTVRAAIEFALRELDVENIIVCGHSRCGACKAYLEASQGKEEQAISRLSEYVSNSESLEKVMANDVLAQIKNLMTFAAAIARRSLDDNKVIAVFHLVTGDMYYSLDGQKFKIVSEDGSDLVE |
| CA26 | MKRLIEGAIQFQRDYFASHPDRFQQLIDEGQNPEVLFITCSDSRIEPAILTASAPGDLFMLRNVGNLIPAYGEMLGGVSSAIEYAVAALQVRDIIVCGHTHCGAIKGLLDTEAIRQIPHVRTWLEYIKPVTDALPTTLDGGDRAPGDEARLRQVVQEHVNLQLKHLLTYPIIKAQLERGKLELLGWHYIIENGEVFAYDPDADTFRAFSS |
| CA27 | MSTDSSTPSEAWKQLEEGNRRFVSGQILHSRNRQDMTASAKEHEDGQHPWAVVLTCSDSRVPPESLFDQRPGDIFVIRTAGQVFSAEIGGSSAIDYAIKALGVTDVIVMGHTSCGAMTATVEAYSQGDPLSTTNLGSALKAIEPAVQQALEESDGLDAEDEAMVRHAVDLNVRRSIESVRARTTVPTALQHGSLKIVGATYHLEDGRTEII |
| CA28 | MSQVPDTPPSGEPSTAAQSERDADDELKYLVDGVRHFKNEVFPGQEELFKSLAKAQHPKTMFITCADSRLLTSEVTSASAGDIFVVRNICNTAAVQVDGGGVTSLTYPIIHSFEVAAAVLEVKHIIVCGHSDCSSMKGAVNPDGLTKFVADYVQNAKPALERFPSEKLNSVAEKNVRSSAELIRRSPMLAEMEEKGELNLHAWVFEIADGKLFFTEEAGEYVPVS |
| LDH1 | MSAIENSKVTVVGAGSVGTAIAYACLIRGVGRHVALYDLAAPKTQAEVLDLNHGLQFVPVGTVEGSDSLAVCADADVIAICAGAKQKPGQSRIELAGVNVEICKGLIPKLLEVAPEALILMVTNPVDILTYAALKFSGLPPHRVIGSGTVLDSSRFRFLIAQHGRVAVSNVHAYIAGEHGDSEIPLWSSANVSGTPLRDFAVPGRGEFSAADEDELFIDTRDAAHTIIEGKGATNFAIGLATTRIIEAVLNDENSVLPVSSLLTDYRGISEVCLSVPSIVDRGGVAARLLPRLSTGELELLQRSAQALRAVASSLGL |
| LDH2 | MITSRKVAVVGCGFVGSSSAYALVNQGVTDEIVLVDLNHQRAEADARDLRHAVPFAHPVKVWSGDYRDLSDCSLAVITAGGAAQPGETRLDLVRRNTAIFKSVVPQVAAANPNGILLVATNPVDILTYVAQRISGLPSSRVIGSGTVLDTARFRYLLGEHFDVDPRHIHAYVVGEHGDSEVATWSLANVAGVPLERFCKLRGVSMDEKEALAAIEDEVRYAAYQIIERKGCTYFAVATSIMRIVKAILHDEYSILPVSAHLDGEYGIEGLYLGVPSVVNRSGVRETLEIPLDGEEQAALEASASAVHGTLERLRDRA |
| LDH3 | MSDRFRPSKLVIVGAGNVGATFAYALLLSGLAAEIILIDKDRARAKGEVMDLTHAVAFLNPTVVIEAGTYDECADAAIVVVTAGAAQKPGETRLELAKTNADIFKTIIRDVMANNTDGILLVVTNPVDVLTYITWKLSGFPSNRVIGSGTVLDTARLKYLLSQHCDVDARNVHGYIIGEHGDSELPWSLASVAGTPIDQYCQSHGITFSEEDRAQIFQDVRDSAAAIIENKGATYYAVAAGLMRITRAILRDENTVFSVSTLIQGYHGIDDIYLSLPTVVNKNGVREVIELELDANERELFQKSVSHLKDIINNLGI |
| LDH4 | MIETRKASKVTVIGAGSVGATFAYALLNSGIATTVVLTDPNERLAEAQVQDLAHAAPIGRPMHISAGGYEDCGGAAVTILTAGAAQKPGETRLELVQKNVSIFKEIIPKVVAANPNGIVLLVATPVDVLTYAAIKFSGLPSGRVIGSGTVLDTSRFRWILGNHFGVDPASVHAYIIGEHGDSEFPAWSVANIAGMRLDDYAAMHKIEGALDPERLEEIFINTRNTAYEIIAAKGATHYAIGLSVARIVECILRNHSVMSVSNLVNGPYGIDGLYIGMPSIVGREGIEGVLELPVEEEELGRLRHSANVLRETIRKIDF |

| | |
|---|---|
| LDH5 | MRRAGASLPRTRVAIVGTGAVGTAIAYACLIRGSADSVALFDVNKAKVEAEALDIAHALPF<br>SHSMDIVGGSDIDVVAGSQIVAITAGSKQRPGQSRLELAATNVDMTRKLIPQLVEQAPQAV<br>VLMVTNPVDVLTYAALRFDPAPEGVLGSGTVLDSGRLRDLLARRAGVSLASVHAMVAGE<br>HGDSEVAVWSSATIGGVPLLDWRGWNGSPDALADALRDEVANAAYEVIAGKGATNYAV<br>GLAATRIVEAILNEEHRVLPVSSVLSGFRGVADVALSLPSVVDAHGVVRVLEVPMTEPERA<br>AIDRSAAALQKVQESLGL |
| LDH6 | MAVPSSADGGLSRRTRSSKVAIVGAGSVGATLAYACLVRGVAKTVALYDLNEAKVRAEVL<br>DLSHGLEFVPQASVVGSEDIDVVEGSHVVAITAGAKQKPGQSRLDLAGANVALTRDLMP<br>QLVAVAPDAILLVVTNPVDVVTYVAQKISGLGPGRVIGSGTVLDTSRLRHLLAERLGVSVQ<br>NVHAVIAGEHGDSEFPLWSTANIGDVPLEQFSIPGPWSTADLAALFEQTKDAAYAVIAGK<br>GATNYAVGLAATRIVEAVLGDENRILTISSLVTDYHGIGGVALSVPSVVGARGLERVLDVP<br>MSEAEEAGLRGSAQTIASIQGSLGF |
| LDH7 | MAVLEKSNKIVIIGAGFVGTSIAYTLALSGLASEIVLVDRNQETAQAQALDLAHGLPFAHPM<br>DIYTGSYEDCHGADIIIVSAGATQKPGESRIGLVKKNIAIFKSIAPQIVNYYHQDAILLVVSNP<br>VDILSYVTLAISGFPRERVIGSGTVLDTARLKALLASHFNIDARNVHTFIIGEHGDSSVPAWS<br>LANISGVPINHFCSTCKICNDKELKNSIFHETKDSAYEIIEKKGSTSYAIALATVDIVEAVLRD<br>QGRILPVSNMVEGVYNLEDIALSIPTVLNEKGISRIIEMPMGQEEVENLKKSAEVLKDTLSS<br>LGL |
| LDH8 | MNEKVNRVALIGAGGVGSSFAYALTAQGVADEIVMIDLDKARADAHALELRHGLRFASPQ<br>RVWAGGYEDCKDADVVVICAGAAQKPGETRLDLVQKNASIFSGIIENIMKAGFNGIFLVAT<br>NPVDILTYATYKLSGFPTHRVIGSGTTLDSARLKTIIGEELQVDPRDVNAYVIGEHGDTELA<br>MWSHATIGGIPVTDYCKLRHPDGGVLTPEDMNQIFINVRDAAYQIIDYKGSTYYGVASALA<br>RITKAIVRDENTILPVSTLLDGQYGLDHVFLGIPSIVNGSGIGQLYKMELSDTEKDRLHQSA<br>EALKAVLAQIGK |
| LDH9 | MPELDRRKVGIVGVGSVGATTAYALLIRGVGAEIVLFDRNARTAEAHAMDLQHGLPFVPP<br>AQTRAGGYGPLAGCGVIILCAGVTQRAGETRLDLLQRNAAVFEEVIPRLAASAPDAVLVVV<br>TNPVDVLTHAALAAAGPDARVIGSGTTLDTARFRALLGGALGVDVGHVHAFVVGEHGDS<br>EVVLWSGASVGGVPLEAFAEQHGRSLAEGDIDRLFANTRRAAYEIIAGKGATYYGVAGIV<br>GRLVEAIARDERRVLTVSARTDAVPGLPPVVSISLPRVVGRDGVVDTLLLPLDDGEREGLV<br>ASARVLESAAAAVPEEESSVPREVRPTG |
| LDH10 | MSEISTVKPRKVAIVGCGFVGCATAYTLMQSGLFNEMVLIDANRKKAEGEALDITHGLPFA<br>RQMDIYAGDYSDCADAAIVIITAGAAQKPGETRLDLVKKNAMIFRSIVPEIARYNPEGILLIV<br>SNPVDVMTHIAVKLSGYSSGRVIGSGTVLDTARLKYLLGEHLGVDSRSVHAFVVSEHGDS<br>EIAAWSSANVSGIPLEEFCAMRGHFDYEKATIDKIADSVKNSAYDIIARKHATYYGIAMSVK<br>RICEAIVRDEKSILPISSLVEGEYGLDDVYIGTPAIVCGEGAEQILELPLNQEEEAKFAASAK<br>SLRDIIDSFSL |
| LDH16 | MSAIENSKLTVVGAGAVGASVAYACLIRGTARHVALYDINAPKVEAEVLDLAHGAQFTGS<br>SDITGGSDISVAEGSHVVVITAGAKQKPGQTRIELAGVNAKIIENLMPKLLELAPDAIYVVVT<br>NPCDVLTVAAQKITGLPPTQIFGSGTVLDTSRLRWILAKRAGVSISSVHAQIIGEHGDTEFP<br>LWAQARVGPVPILDWVPTDGEKPFTAEVRDDIAHEVVNAAYKVIAGKGATNYAIGLSSARI<br>VEAIIGDEHAVMPVSNVLNDWHGISDVALSVPSIVGREGVVNRLPLPLSADEESALKASAD<br>ALRETARSLGF |
| LDH17 | MTEKQRKKVILVGDGAVGSSYAFALVNQGIAQELGIVDIFKEKTQGDAEDLSHALAFTSPK<br>KIYSAEYSDCHDADLVVLTSGAPQKPGETRLDLVEKNLRINKEVVTQIVASGFKGIFLVAAN<br>PVDVLTYATWKFSGFPKERVIGSGTSLDSARFRQALAEKIGVDARSVHAYIMGEHGDSEF<br>AVWSHANVAGVGLYDWLQANRDVDEQGLVDLFVSVRDAAYSIINKKGATFYGVAVALARI<br>TKAILDDEHAVFPVSVYQEGQYEVFIGQPAIVGARGIVRPVNIPLNDAELQKLQASAKQLQ<br>DILDEAFAKEEFASAVK |
| LDH18 | MSSVLQKLITPIASGPAEPPRNKVTVVGVGQVGMACAVSILLRDLADELALVDVMEDKLK<br>GEMMDLQHGSLFLKTSKIVADKDYAVTAHSRIVVVTAGVRQQEGESRLNLVQRNVNVFK<br>C<br>IIPQIVKYSPNCTILVVSNPVDVLTYVTWKLSGLPRHRVIGSGTNLDSARFRYLMAERLGIH<br>STSFNGWILGEHGDTSVPVWSGANVAGVSLQTLNPDIGTDGDKEHWKATHKAVVDSAY<br>EVIRLKGYTNWAIGLSVADLTESLVRNMSSVSAVSTSVKGMYGIDNDVFLSLPCVLNSSG<br>VASVVNMTVTDEEIAQLQKSAETLWGVQRDIKDL |

| | |
|---|---|
| LDH19 | METSKVVILGASRGQTESYLFTRLCKDNPSIDAEFVGFGLETDLDAFVNSLSDLGATLDSLLTKAGATSPKTITASSVKASVAEKVGRIDFLIYPLTSFISAGSDLAVVDSSLLATSLGIGFSLVEEKIKELENRGARAATIYSSFTHEVSKTKLLLGEIVPAILDLPTVVDTTELEEITRLLKNRIESAAGLPVVGKISEEISKKAEEAASELSIPLELLSVLSQIGKEKVSLVEKSASASILGAKILAVLLDVYAILPLSVIDKGGLEDTFYSPEESIEDLGAEVVAPVLAGGEIPVKLGEDGIKRVFAEAEFERFEKAAEEVKNS |
| LDH20 | MTSTKQHKKVILVGDGAVGSSYAFALVNQGIAQELGIIEIPQLFEKAVGDALDLSHALAFTSPKKLYAAQYSDCADADLVVITAGAPQKPGETRLDLVGKNLAINKSIVTQIVESGFKGIFLVAANPVDVLTYSTWKFSGFPKERVIGSGTSLDSARFRQALAEKLDVDARSVHAYIMGEHGDSEFAVWSHANIAGVNLEEFLKDTQNVQEAELIALFEGVRDAAYTIINKKGATYYEIAVAMARITKAILDDENAVLPLSVLVREGQYGVKNVFIGQPAVVGANGVEEIIELPLNDAERQKMAASAKELQAIIDEAWKKESE |
| LDH21 | MNTFEPSRVAIVGSGAVGTSLAYAALTRGTAQEVVLYDIDGKRVRAEALDLQHAVPFTGGMRVEGGSDIDIAAGAHVVVVTAGAKQAPGQTRLDLAGVNVGILQSLLPQVQAARPDAVIMLVTNPCDVLTVLAQTSTGLPPHRVFSSGTVLDTSRLRWLLAAEAGVAIKNVHAYVAGEHGDTEFAAWSTATIGGVPLLDWSTPGHPVFTAEQLARIQDAVVHSAYRVIEGKGATNYAIGLAATRIVEAILRDQDVVLPVSTVLDDYRGVSGVSLSLPTVVSSAGALTVSETPLSVQEEARLADCARTLRATADSVRP |
| LDH22 | MTKINRVALIGSGFVGSSYAFALINQGIANELVLIDVNKEKAEGDAMDLNHGKAFAPQPTTIWAGNYHDCNDADIAVICAGANQKPGETRLDLVEKNLRIFKSIVQEVMGSGFDGIFLVATNPVDILTYTTWKFSGLPKDRVIGSGTILDTARFRYLLGEYFDVDTRNVHAYIIGEHGDTELPVLSSANVGGVPVSKIAKQLENNEDYDAEDLEDIFIRVRDAAYHIIDRKGATYYGIAMGLARITRAILHNENAVLTVSAYLEGEYGQNDVYIGVPAVVNRSGIREVVELELNDYELHHLNDVYQNTREILTTVF |
| LDH23 | MTATKQHKKVILVGDGAVGSSYAFALVNQGIAQELGIIEIPQLFDKAVGDALDLSHALAFTSPKKIYAAQYSDCADADLVVITAGAPQKPGETRLDLVGKNLAINKSIVTQVVESGFNGIFLVAANPVDVLTYSTWKFSGFPKERVIGSGTSLDSARFRQALAEKIGVDARSVHAYIMGEHGDSEAVWSHANVAGVKLEQWLQANRDLNEEGLVELFVSVRDAAYSIINKKGATYYGIAVALARITKAILDDENAVLPLSVFQEGQYGVNNVFIGQPAIVGAQGIVRPVNIPLNDAETQKMQASAKELQAIIDEAWKNPEFQEASKN |
| LDH24 | MATLKDQFKPPAPSAVPNNKITVVGVGQVGMACAISILGKSLSDELALVDVLEDKLKGEMMDLQHGSLFLRTPKIVADKDYSVTANSKIVVVTAGVRQQEGESRLNLVQRNVNIFKFIIPQIVKYSPACIIIVVSNPVDILTYVTWKLSGLPKHRVIGSGCNLDSARFRHLMAEKLGIHASSFNGWILGEHGDSSVPVWSGVNVAGVSLQNLNPDMGTDNDSENWKVVHETVVKYGYEVIKLKGTNWAIGLSVADQIEKLINNLSLVDEVMTEGVTDESGSSVKKVIEIPLVLGNNGIGDVMLTVPAILTRNLNVSRVVKNVKTIKKVFQKIIF |
| LDH25 | MKKIVLIGDGAVGSTYAFALMQQRICEEIVIINRSKNKAEAIDLSHGMPFTEQTKVYSGAYEDARDSDIVIITAGAPQKVGQSRLDLMAINAKIAQSIVDSIMASGFDGYFVIASNPVDILSYAWQFSGLPKERVIGSGTSLDTARLRVQLSKHFSVSSSSVDSFILGEHGDTDFAVWSHANIGGRPLMQYLEETKKVKEKDLVEMSQQVKNAAYEIIKSKGSTYYGIGVALDRITNAILGNNHRILAVSSLIKNQYDLSHNIYIGVPAVIGANGIDKQLEIDFTANEKKKLTLSAKTLKEAQTKLNS |

**Table S7. LDH activity data at different pH concentrations**

| Testing ID | pH 4.5 | pH 7 | pH 9.5 |
|---|---|---|---|
| LDH1 | 0.27 ± 0.16 | 0.19 ± 0.15 | |
| LDH2 | 0.26±0.19 | 0 | 0 |
| LDH6 | 0.55±0.05 | 0.19±0.13 | 0.13±0.03 |
| LDH7 | 0.61±0.19 | 0.92±0.30 | 0 |
| LDH8 | 0 | 0.44±0.23 | 0.22±0.11 |
| LDH9 | 0.24±0.19 | 0 | 0 |
| LDH10 | 0.36±0.16 | 0 | 0.12±0.05 |
| LDH16 | 0.42±0.10 | 0 | 0.16±0.11 |
| LDH18 | 0 | 0.32±0.13 | 1.50±0.43 |
| LDH19 | 1.91±0.25 | 0.94±0.16 | 0.35±0.05 |
| LDH20 | 0.61±0.11 | 0 | 0.14±0.03 |
| LDH21 | 1.85±0.31 | 0.78±0.55 | 0.41±0.03 |
| LDH23 | 0.32±0.14 | 0 | 0 |
| LDH24 | 1.52±0.30 | 0.72±0.33 | 0.19±0.07 |
| LDH25 | 0.49±0.06 | 0.31±0.28 | 0.14±0.03 |
| Control1 | 0.42±0.3 | 0 | 0.08±0.02 |
| Control2 | 0.62±0.29 | 0.23±0.19 | 0.11±0.05 |
| Control3 | 0.24±0.23 | 0 | 0 |

**Table S8: Conditions used for experimental protocol of cascade 2**

| Phase | Rate (°C /min) | Oven temperature | Hold time(min) | Run time(min) |
|---|---|---|---|---|
| Initial | 0 | 40 | 0.5 | 0.5 |
| Ramp up | 120 | 300 | 1.5 | 4.167 |

**Table S9:Reaction conditions for one-pot enzyme cascades:** using LDH18 and LDH19 along with proprietary transaminases (RTA-25, STA-14, RTA-57) and glutamate dehydrogenases (GDH-101) from Johnson Matthey. Cascade 1 utilises GDHs for gluconic acid to glucose conversion. Cascade 2 employs transaminases for alanine to pyruvate conversion coupled with LDH conversion of pyruvate to lactate. Experiment: naming scheme used for each experiment. RTA and STA specify R-selective and S-selective transaminases, respectively. T and pH list the optimised temperature and pH conditions based on the enzymes involved in each experiment.

| Experiment | Transaminase | LDH | GDH | T (°C) | pH |
|---|---|---|---|---|---|
| LDH18 - GDH-101 pH 9.5 | - | LDH18 | GDH-101 | 45 °C | 9.5 |
| LDH19-GHD-101 pH 5 | - | LDH19 | GDH-101 | 45 °C | 5 |
| RTA-25-LDH18-GDH-101 pH 9.5 | RTA-25 | LDH18 | GDH-101 | 45 °C | 9.5 |
| STA-14-LDH18-GDH-101 pH 9.5 | STA-14 | LDH18 | GDH-101 | 45 °C | 9.5 |
| RTA-57-LDH19-GDH-101 pH 5 | RTA-57 | LDH19 | GDH-101 | 45 °C | 5 |

**Table S10**: **Evaluation of LDH18 and LDH19 concentration pre and post-lyophilisation using the Bradford assay**

| **Experiment** | **Protein concentration (obtained with Bradford assay)** |
|---|---|
| LDH18 pre-lyo | 13.7 $mg_{protein}$/mL |
| LDH18 post-lyo (prepared 10 $mg_{powder}$ /mL) | 4.1 $mg_{protein}$/mL |
| LDH18 post-lyo (prepared 1 $mg_{powder}$ /mL) | 0.4 $mg_{protein}$ /mL |
| LDH19 pre-lyo | 14.4 $mg_{protein}$/mL |
| LDH19 post-lyo (prepared 10 $mg_{powder}$ /mL) | 4.9 $mg_{protein}$/mL |
| LDH19 post-lyo (prepared 1 $mg_{powder}$/mL) | 0.5 $mg_{protein}$/mL |

### Supporting Information References


1. Dill, K. A., Ghosh, K. & Schmit, J. D. Physical limits of cells and proteomes. *Proc. Natl. Acad. Sci. U. S. A.* **108,** 17876–17882 (2011).
2. Holehouse, A. S., Das, R. K., Ahad, J. N., Richardson, M. O. G. & Pappu, R. V. CIDER: Resources to Analyze Sequence-Ensemble Relationships of Intrinsically Disordered Proteins. *Biophys. J.* **112,** 16–21 (2017).
